# Dissecting Small Noncoding RNA Landscapes in Mouse Preimplantation Embryos and Human Blastoids for Modeling Early Human Embryogenesis

**DOI:** 10.1101/2025.10.07.680944

**Authors:** Savana Biondic, Zhao Cheng, Richard Yin, Thorold Theunissen, Sophie Petropoulos

**Author notes:** These authors contributed equally (Savana Biondic, Zhao Cheng).

## Abstract

Small non-coding RNAs (sncRNA) exert regulatory functions in mammalian cells; however, their expression dynamics and contribution during preimplantation development are not entirely understood. In this study, we apply Small-Seq to generate a comprehensive single-cell atlas of sncRNA expression in mouse oocytes, sperm, and embryos (2-cell to 64-cell stages), and compare these dynamics with human embryos at equivalent stages. In both species, all sncRNA subtypes are expressed, but only microRNAs (miRNAs) and small nucleolar RNAs (snoRNAs) display cell type-specific patterns. In mice and humans, miRNAs and snoRNAs from the *Dlk1-Dio3* locus were upregulated in the inner cell mass (ICM). Human trophectoderm (TE) is enriched with primate-specific C19MC miRNAs, which show relatively low expression in the ICM. In contrast, the mouse lacks a TE-specific hotspot, as C2MC miRNAs are higher in the ICM. Nonetheless, differentiation-associated miRNAs (e.g., miR-24-3p, miR-200c-3p) are high in the TE of both species. Further, we profile the sncRNAs of human blastoids (stem-cell-based embryo model) and determine that they largely recapitulate the human blastocyst. We envision broad utility of this dataset as a resource for future studies seeking to dissect the functions of individual sncRNAs in early development, and for advancing applications in stem cell-based embryo models and assisted reproductive technologies.

## Introduction

The preimplantation period of mammalian development is critical for ensuring embryo competence and shaping the overall trajectory of an organism’s growth. During this time, embryonic genome activation (EGA), cell divisions and a meticulously orchestrated series of molecular events transform a fertilized egg into the blastocyst. In addition, previously totipotent cells of the embryo begin to undergo lineage specification, giving rise to the first distinct cell types: the trophectoderm (TE; prospective placenta) and inner cell mass (ICM), the latter subsequently specifying further into the epiblast (EPI; prospective embryo proper) and the primitive endoderm (PrE; prospective yolk sac)^1^. With these specialized cell types, the blastocyst is able to implant in the uterine wall, a necessity for establishing a viable pregnancy. Single-cell genomics have been indispensable in advancing our current understanding of the molecular mechanisms underlying mammalian preimplantation development and lineage specification. Single-cell RNA sequencing (scRNA-seq) for instance has provided detailed maps of the dynamic transcriptomic patterns throughout early development, and has illuminated the roles of key transcription factors, signaling pathways, and other mechanisms that influence pluripotency, plasticity, and embryonic cell fate decisions^2–10^. In addition to protein-coding transcripts, cells possess a group of endogenous RNA molecules typically less than 300 nucleotides in length known as small non-coding RNAs (sncRNAs)^11,12^. Once deemed non-functional artifacts due to their lack of protein-coding capacity, sncRNAs are now recognized as intricate cellular players, exerting their influence on multiple levels, notably in the realm of epigenetic gene expression regulation^11–13^. Given the growing awareness of the regulatory roles executed by sncRNAs, there exists a compelling rationale to delve into their potential contributions to preimplantation development.

With the advent of single-cell genomics and increased access to human embryos, our understanding of human embryogenesis has increased over the last few years. However, research with human embryos is often faced with limitations related to access and is governed by stringent ethical and legal legislation. As such, we rely heavily on modeling systems. The gold standard model currently utilized for translation to human preimplantation development is the mouse, which shares some conserved mechanisms with the human but differs in timing of milestone events and expression dynamics of some key transcription factors and signaling pathways involved in lineage specification, commitment and pluripotency^1^. While we and others have recently profiled sncRNAs throughout human preimplantation development yielding numerous valuable insights^14,15^, the same has not been done for the commonly used mouse model. Given the well-characterized molecular differences in mouse and human preimplantation development^1^, a cross-species comparison of sncRNA expression dynamics is critical for assessing the translatability and inferential validity of mouse sncRNA studies to human embryogenesis. Similarly, the expression patterns of sncRNAs in the recently developed human embryonic stem cell (hESC)-derived blastoid models remains unknown^16–22^ and would provide an additional molecular layer for benchmarking, providing further insights into the utility of human blastoids as a tool to better understand human preimplantation development.

A comprehensive single-cell delineation of sncRNA expression in mouse germ cells (sperm and oocytes) and throughout mouse preimplantation embryonic development (preceding and during lineage specification) would help deepen our understanding of the global gene regulatory landscape in the preimplantation embryo and drive further research into the functional roles of sncRNAs (such as miRNAs). Utilizing a combined approach with Small-Seq^23^ and our method, Co-Seq^24^, we provide a resource characterizing sncRNA dynamics throughout mouse preimplantation development, identifying miRNAs as important potential regulators of various developmental processes such as the first cell fate decisions. We also perform a cross-species comparison with our previously published human single-cell Small-Seq dataset, and our new human blastoid Small-Seq data generated in collaboration with the Theunissen Lab^16^, to identify how well these models recapitulate human preimplantation development. This information could be leveraged to further improve stem cell models like the blastoid, and/or could pave the way for the development of non-invasive embryo selection methods such as sncRNA-profiling of spent culture media, to overall enhance the efficacy of assisted reproductive technologies.

## Results

### Overview of sncRNA expression throughout mouse preimplantation development

In order to comprehensively delineate sncRNA expression throughout mouse preimplantation development, embryos spanning the 2-cell to 64-cell (blastocyst) stages were collected and dissociated into single cells for profiling with Small-Seq (Fig. 1a), fully encompassing the processes of compaction and the first lineage specification. In addition, MII oocytes and sperm were collected and profiled to potentially provide insight into sncRNA inheritance from germ cells to the early embryo (Fig. 1a). After quality control (QC) (Extended Data Fig. 1a,b), a total of 680 cells isolated from 57 different embryos were analyzed (Fig. 1a and Supplementary Data 1). In addition, 27 oocytes and 16 bulk sperm samples (from 3 different male mice) were analyzed (Fig. 1a). In the resulting data, read length distributions from each sncRNA subtype show expected patterns, with primary peaks at 21-24 nt for miRNAs^23,25,26^, 18-30 nt for piRNAs^26–29^, 65+ nt for snoRNAs^14^, and 30-41 nts for tRNA-derived small-RNAs (tsRNAs, particularly 5′ and 3′ tRNA halves)^23,30,31^ (Fig. 1b,c), verifying the quality of data obtained. piRNA length distribution shifts between mouse sperm, oocytes, and preimplantation embryos, with a 1U nucleotide bias at the 5′ end confirming their piRNA identity (Fig. 1c and Extended Data Fig. 1d).

**Fig. 1.**
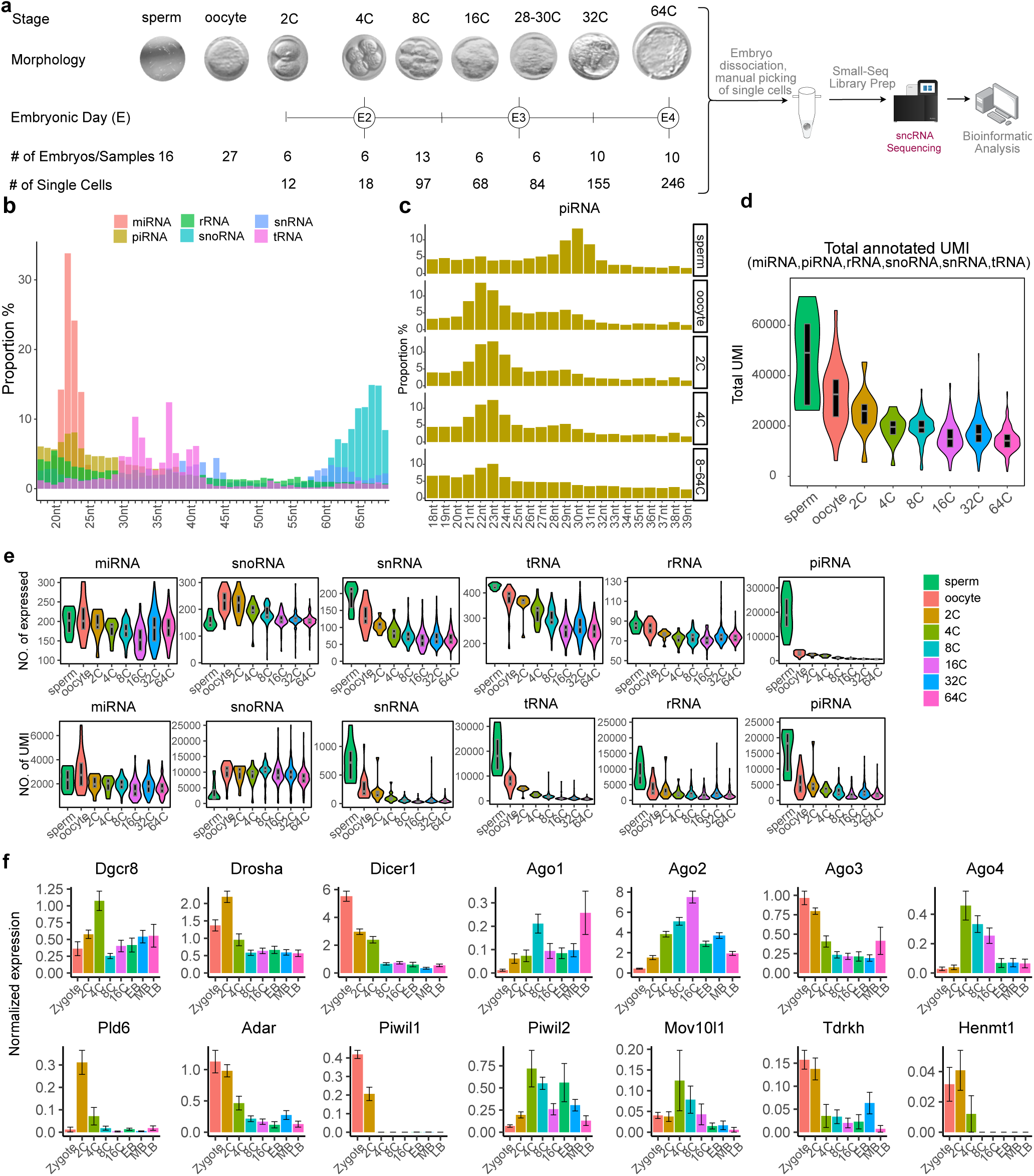
Single-cell profiling of sncRNAs in mouse embryos. **a,** Overview of the experimental workflow, including the total number of cells and embryos analyzed at each stage of mouse preimplantation development. **b,** Percentage of total annotated unique molecular identifiers (UMI) by sncRNA subtype of various sequence lengths (nt = nucleotides). **c,** Percentage of UMI sequence lengths distribution from piRNA stratified by day of embryonic development. **d**, Total UMIs distribution by day of embryonic development. **e,** Total number of expressed sncRNA subtypes and UMIs throughout mouse preimplantation development. **f,** RNA expression of sncRNA biogenesis machinery members from re-analysis of Deng et al., 2014 single-cell RNA sequencing dataset. 2C = 2-cell, 4C = 4-cell, etc.

The total number of unique molecular identifiers (UMIs) detected gradually decreased as development progressed (Fig. 1d). While the number of detected miRNAs, snoRNAs, and rRNAs remained fairly consistent, there was a decrease in the number of tRNAs and piRNAs with developmental time, suggesting the majority may be inherited from the sperm and/or oocyte (Fig. 1e and Extended Data Fig. 1c). These patterns are very similar to what we recently outlined in human preimplantation embryos^14^, and to what has previously been found in whole mouse embryos^27,32^. Notably, most sncRNA subtypes exhibited discernible autosomal chromosome distribution patterns, with preferential enrichment found on specific autosomes (Extended Data Fig. 1e). In addition, sncRNAs in sperm, oocytes and embryonic cells had different contributions from and distributions in the mitochondrial DNA (Extended Data Fig. 2). 21.1% of sperm UMIs were derived specifically from mitochondrial DNA, whereas only 4.9% and 5.3% mapped to the mitochondrial DNA in oocytes and 2-cell embryos respectively (Extended Data Fig. 2a), consistent with observations from previous studies^29^. This proportion decreased further with developmental progression, from 4.8% in 4-cell embryos to a mere 2.14% in 64-cell blastocysts (Extended Data Fig. 2). Detailed analysis of these mitochondrial DNA reads indicated that these mitochondrial contribution discrepancies mainly result from 2 regions, and are mostly tRNA- and piRNA-related (Extended Data Fig. 2b,c).

We next aimed to better understand the change in expression patterns observed for the sncRNA biotypes by determining the expression of key components involved in their biogenesis. To do this, we re-analyzed the previously published Deng et al., 2014 single-cell RNA sequencing dataset of mouse preimplantation development^10^. We observed a general decrease in the mRNA expression of members of the microprocessor complex (*Drosha, Dgcr8*) as well as the pre-miRNA processing enzyme *Dicer1* as development progresses (Fig. 1d), in accordance with prior studies^33,34^. The expression of the Argonaute (*Ago1-4*) proteins, which form the miRNA-induced silencing complex (miRISC) with mature miRNAs to regulate the expression of target transcripts, varies depending on the paralog. While *Ago2* expression peaks at the 16-cell stage to then decrease in blastocysts, *Ago3* and *Ago4* peak in 2- to 4-cell embryos and steadily decrease as development progresses (Fig. 1f). Also of note, expression of major players involved in piRNA biogenesis and maturation, including *Pld6, Piwil1*, *Mov10l1, Tdrkh* and others^35^, underwent steep decreases in expression after the 2- to 4-cell stage (Fig. 1f), possibly explaining the decrease in piRNA expression associated with developmental progression.

### Identification of cell-type specific miRNA profiles

Previous studies, including works from our lab and others, have revealed dynamic changes in miRNA expression across developmental stages and lineages in human preimplantation embryos^14,15^. However, miRNA expression at the single-cell level in the corresponding mouse embryo model remains largely unexplored, and we sought to determine whether we could similarly identify miRNA signatures associated with the mouse ICM and TE. To circumvent the lack of known defined miRNAs signatures for ICM and TE in mouse, we leveraged our previously established method of single cell mRNA-sncRNA Co-sequencing (Co-Seq)^25^ on 313 cells (post-QC) from 39 embryos spanning the 2-cell to 64-cell (blastocyst) stages (Supplementary Data 1). This technique enables parallel sequencing of the transcriptome (with Smart-Seq2) and the sncRNAome (with Small-Seq) within the same cell, thereby facilitating the characterization of sncRNAome features in cells with lineages accurately resolved from transcriptomic data^14,25^.Furthermore, it establishes a data framework for systematic investigation of gene–sncRNA expression relationships.

To assess Co-Seq data quality, we confirmed that quality control metrics such as the number of sequenced sncRNA and mRNA reads and the diversity of expressed miRNAs and genes were comparable to those from whole cells (Extended Data Fig. 1a,b). We then integrated the mRNA portion of the Co-Seq data with a previously published Smart-Seq2 dataset spanning mouse preimplantation development to identify cell types (Fig. 2a,b)^36^. Blastocyst cell identities (ICM and TE) were assigned following data integration (Extended Data Fig. 3a,b) which was further supported by expression of known lineage markers (e.g., *Sox2* and *Nanog* (ICM), *Dppa1* and *Gata3* (TE)) (Extended Data Fig. 3c). Subsequently, cells of known lineage were indicated onto a Uniform Manifold Approximation and Projection (UMAP) constructed from the miRNA expression profiles of the sncRNA portion of the Co-Seq data (Fig. 2a). Similar to observations in humans^14^, the miRNA expression patterns of cells organized non-randomly into two distinct clusters corresponding to ICM and TE from 16- to 64-cell stages (Fig. 2a). After confirming that lineage could be reliably distinguished by miRNA expression and identifying lineage-marker miRNAs (Fig. 2a and Extended Data Fig. 3d), we integrated our Co-Seq data with full-cell Small-Seq data, thereby assigning lineage annotations to the Small-Seq dataset (Fig. 2b,c and Extended Data Fig. 3e) and determined the top differentially expressed miRNAs across developmental stage and cell types (Fig. 2d and Supplementary Data 2).

**Fig. 2.**
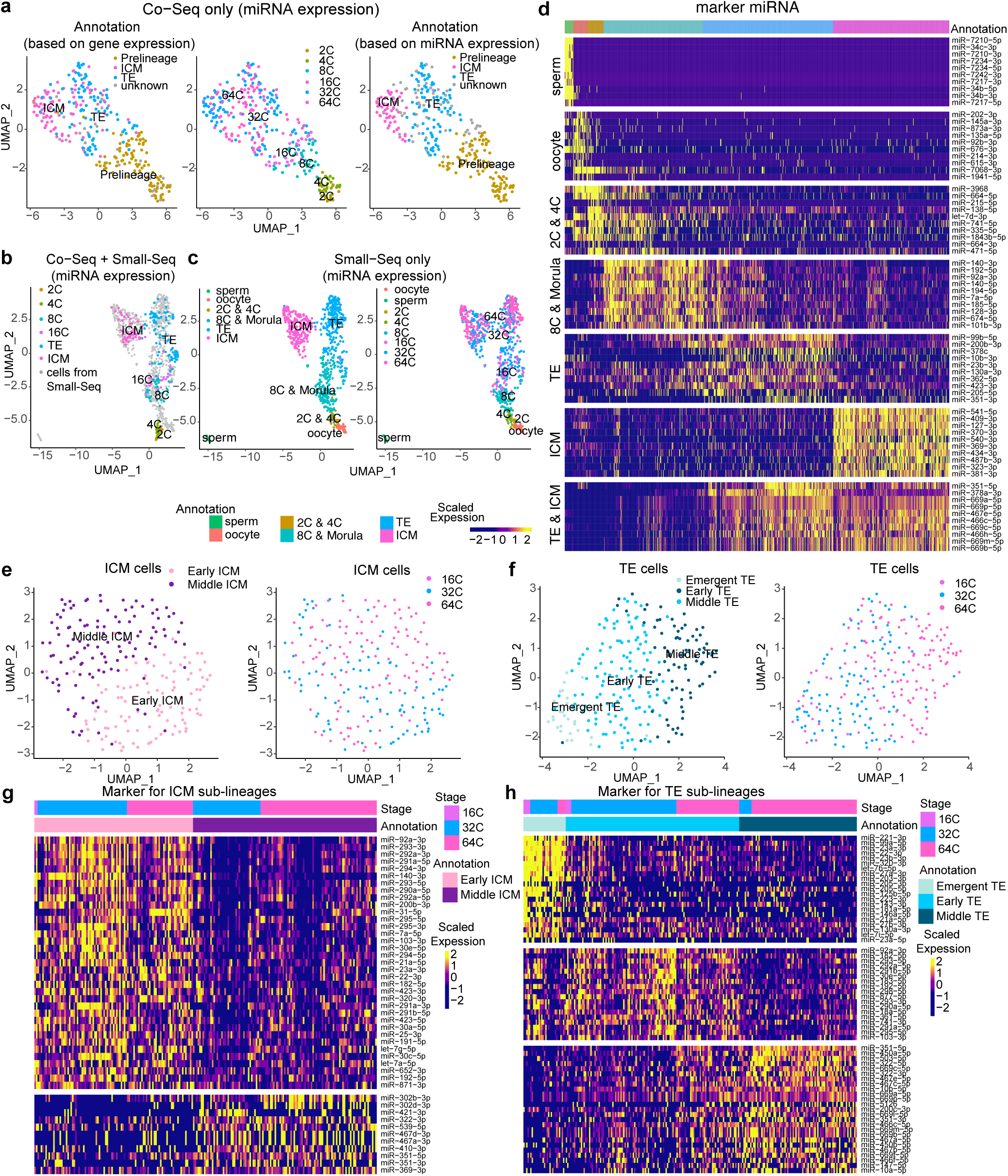
Lineage identification and miRNA dynamics across mouse embryo development. **a,** Uniform Manifold Approximation and Projection (UMAP) showing the dimensional reduction of Co-Seq data (Small-Seq portion) based on miRNA expression. Cells are coloured by annotations derived from gene expression, developmental timepoint, and the final assignment based on miRNA expression. **b,** UMAP showing the integration of cells of known identity (sequenced with Co-Seq) with unknown full cells of the main dataset (sequenced with Small-Seq only) based on miRNA expression. **c,** UMAP depicting only the main dataset (Full cell Small-Seq) from the integration in (**b**) colored by developmental stage and lineage. **d,** Heatmap of the top 10 significantly enriched miRNAs in each lineage throughout mouse preimplantation development. UMAP showing further sub-clustering of miRNA expression profiles among **e,** ICM cells and **f,** TE cells in the full cell Small-Seq dataset. Heatmaps of significantly differentially expressed miRNAs between **g,** the early and middle ICM sub-clusters and **h,** the emergent, early and middle TE clusters.

miRNAs that were highly expressed specifically in the early stages of mouse preimplantation development include specific members of the let-7 family, miR-98-5p, and miR-100-5p (Fig. 2d). Notably, several miRNAs exclusively upregulated at the 8-cell stage (e.g., miR-92a-3p, miR-140-3p, miR-185-5p) (Fig. 2d and Supplementary Data 2) have been shown to regulate Hippo signaling, PI3K/AKT signaling, and E-cadherin–mediated cell–cell adhesion in cancer studies^37–39^, raising the possibility that miRNAs may contribute to the regulation of polarization and compaction into a morula. Further analysis of the ICM and TE miRNA profile clusters revealed sub-clustering within each of the lineages. The ICM could be divided into an early ICM (comprising a few cells from 16-cell embryos, but mostly 32-cell (early) blastocysts) and a middle ICM (consisting primarily of cells from 64-cell (middle) blastocysts), likely reflecting the maturation of the ICM into the EPI (Fig. 2e,g). Examining individual miRNAs associated with each sub-cluster, we identified higher expression of miRNAs from the miR-290-295 cluster, the miR-17-92 cluster, and miR-182-5p—all known to be characteristic of naïve mESCs—in the early ICM^40,41^. The middle ICM was enriched for members of the miR-302 family as well as miR-421-3p, miR-539-5p, and miR-410-3p, which are more characteristic of primed mouse epiblast stem cells (mEpiSCs)^40^ (Fig. 3g). These results indicate that the transition from ICM to EPI is accompanied by a shift in miRNA profiles similar to those observed during the naïve-to-primed state transition in mESCs, suggesting that expression of specific miRNAs may also be important drivers of the pre-vs post-implantation EPI maturation in mouse embryo. Notably, a distinct PE miRNA signature was not identified in 64-cell blastocysts, when a PE transcriptomic signature is known to be present^1,42^. It is possible that the PE may not form a distinct cluster because it is highly similar to the ICM from which it differentiates or the EPI. Alternatively, the PE signature may be masked within the TE, as it has been shown using qPCR that XEN cells (a PE model) have a similar miRNA profile to trophoblast stem cells (TSCs)^43^. Profiling later-stage embryos, though beyond the scope of this study, could help clarify when an endoderm-specific miRNA profile first emerges.

**Fig. 3.**
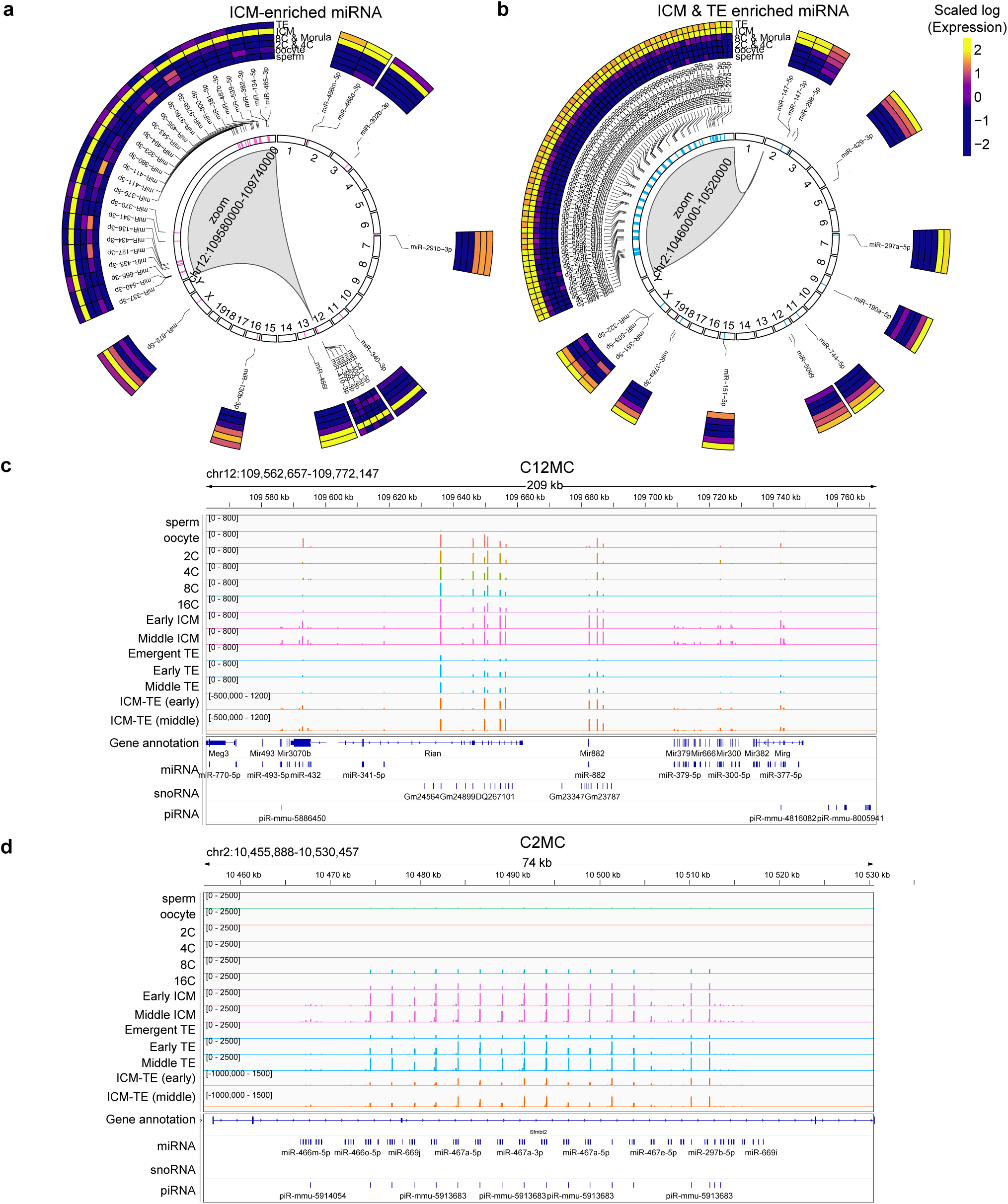
Genomic distribution of ICM- and TE-enriched miRNAs in mouse blastocysts. Circular plots mapping the chromosome localization of **a,** ICM-(zoomed in on chromosome 12 miRNA cluster (C12MC)) and **b,** ICM and TE-enriched (zoomed in on chromosome 2 miRNA cluster (C2MC)) miRNAs. Integrative Genomics Viewer (IGV) showing the genomic coordinates and normalized reads coverage (coloured bars) of miRNAs originating from **c,** C12MC and **d,** C2MC, stratified by mouse embryonic cell types and developmental stages.

The TE miRNA profile could be further divided into three sub-clusters, which we labelled as emergent TE (earliest, contains morula cells), early TE (predominantly 32-cell blastocysts) and middle TE (predominantly 64-cell blastocysts) based on the developmental stage of the source embryos (Fig. 2f,h). The emergent TE had a very distinct miRNA signature characterized by high expression of miR-221-3p, miR-21 and let-7 family miRNAs, which are known to promote differentiation by suppressing pluripotency factors and characterize TE cells^44–49^ (Fig. 2h). The early TE was enriched with miR-290-295, miR-17-92, and miR-183-182 cluster miRNAs (Fig. 2h), some of which have been found to potentiate TS cell self-renewal by targeting cell cycle repressors^50^. The middle TE was defined by increased expression of miR-322 and miR-467, which can reprogram mESCs into a TE phenotype^51^, together with other miR-322 cluster members (e.g., miR-503, miR-351, miR-450a, miR-450b), which are known to increase in differentiated trophoblast cells to target cell cycle activators^50^ (Fig. 2h). Consistent with these observations, studies have shown that TSC differentiation is accompanied by downregulation of the miR-290 cluster and upregulation of the miR-322 cluster^50^. Notably, members of the *Sfmbt2* cluster (e.g., miR-466, miR-467 and miR-669), which are highly expressed in mouse placenta/TSCs^52^, were also upregulated in the middle TE compared with earlier stages (Fig. 2h). The presence of these three distinct miRNA expression profiles across TE development suggests that coordinated changes in miRNA expression may contribute to proper TE maturation, although further studies are needed to determine the precise functions of the different miRNAs at each stage.

### Identification of ICM- and TE-enriched miRNA genomic hotspots

Next we performed chromosome enrichment analysis for miRNA makers in the ICM and TE lineages to determine their genomic origins (Fig. 3a,b). Strikingly, we observed that 79.5% of ICM-enriched miRNAs originate from a region on chromosome 12 (C12MC) commonly referred to as the *Dlk1-Dio3* locus^53^ (Fig. 3a). One of the largest imprinted clusters in mammals, the site consists of protein coding genes preferentially expressed from the paternal allele, and non-coding RNAs (4 long non-coding (lnc)RNAs, Gtl2 (Meg3), anti-Rtl1, Rian (Meg8), and Mirg) expressed from the maternal allele^53^. Enrichment of miRNAs from the *Dlk1-Dio3* locus in the ICM was expected, as they have been shown to be important for ground-state pluripotency in stem cells^54–57^, and are consistently higher in naïve mESCs vs. primed mEpiSCs^58^. In human embryos, we previously found the primate-specific equivalent of this miRNA cluster, located on chromosome 14 (C14MC), to turn on at embryonic day (E)5 and remain highly enriched in the ICM of blastocysts^14^. However, comparative analysis between the species revealed a difference in the developmental expression patterns of this miRNA genomic hotspot, whereby in the mouse 48 out of 96 C12MC miRNAs were expressed in the oocyte and at earlier stages of mouse preimplantation development (mainly within and between the Rian and Mirg regions, Fig. 3c), while an additional 13 miRNAs (e.g., miR-376a-5p and miR-377-5p) are turned on exclusively in the early blastocyst ICM. In contrast, the majority of miRNAs from C14MC in the human embryo are expressed upon emergence of the ICM at ∼E5^14^. Overall, our results suggest that miRNAs from the *Dlk1-Dio3* locus may function in maintaining naïve pluripotency in the mammalian preimplantation embryo like in stem cells, but with nuanced species-specific differences.

The second major miRNA hotpot we identified was the rodent-specific miRNA cluster within the *Sfmbt2* gene (located on chromosome 2, C2MC). As this cluster is known to be essential for placental/TSC development^52^, we postulated it could be important for TE specification in the preimplantation embryo. However, we observed that C2MC miRNAs emerge beginning at the 8-cell stage, and are subsequently upregulated in both the TE and ICM lineages (Fig. 3b,d). The lack of TE-exclusive enrichment suggests that those miRNAs may not act as drivers specifically of TE, but rather contribute to both the ICM and TE. In human blastocysts and stem cells, the primate-specific counterpart, C19MC, is highly expressed in and has important functions relating to the acquisition and maintenance of TE identity (C19MC)^14,59^. However, we find here that a similar TE chromosome hotspot may not be present in the mouse, representing a key potential difference between the species. Whether the mouse possesses a functional equivalent to C19MC, or if it compensates for the absence of a TE-restricted miRNA cluster through other mechanisms, remains to be determined.

### Trajectory and pseudotime analysis of miRNA expression through mouse preimplantation development

Next, we sought to analyze how miRNA expression evolves temporally throughout mouse preimplantation development at a higher resolution. Oocytes and sperm were excluded from the pseudotime analysis to avoid skewing the results, as our primary focus was on constructing trajectories for the ICM and TE lineages. While between 2-cell and 4-cell embryos minimal changes were observed in miRNA profile (only 2 significantly differentially expressed (DE) miRNAs with log2 fold change more than 0.1 and fdr less than 0.05) (Fig. 4a and Supplementary Data 3), substantial differences were observed between subsequent stage transitions (Fig. 4a). The largest changes in miRNA expression occurred during the first lineage specification, where 163 DE miRNAs were identified between the 16-cell morula and the early ICM, and 75 DE miRNAs between the 16-cell morula and the emergent TE (Fig. 4a). In addition, both the ICM and TE populations further underwent considerable changes in miRNA expression between the earliest and latest timepoints analyzed (Fig 4a), suggesting specific miRNAs may be important for the maturation of both cell types. 13 miRNAs from the C12MC cluster showed increased expression during ICM maturation, indicating their potential role in promoting ICM progression.

**Fig. 4.**
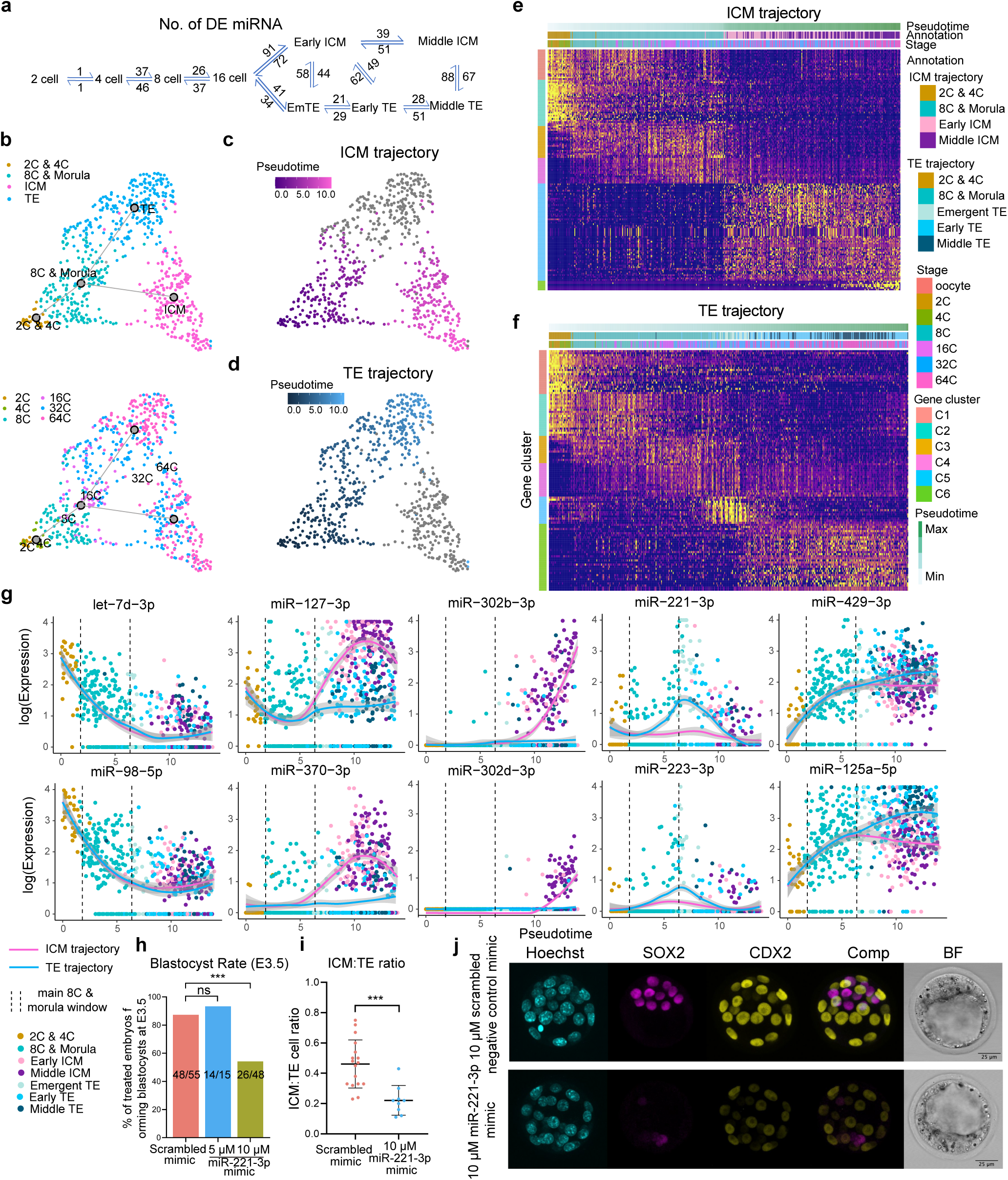
Pseudotime analysis of miRNA expression in early mouse embryonic development. **a,** Differential miRNA expression between developmental stages and lineages. **b-d,** Developmental- and lineage-specific pseudotime trajectories. **e,f,** Heatmaps of lineage trajectory-associated miRNA expression. **g,** Expression of several key miRNAs across pseudotime. The confidence interval (error bands, 95%) is indicated by bandwidth. The measure of centre and confidence intervals were calculated using the “loess” function with default parameters in R software. **h,** Percentage of total embryos treated with scrambled negative control or miR-221-3p mimic at the 2-cell stage that developed into blastocysts by embryonic day (E)3.5. Statistical analysis was conducted using the Chi-Square Test. Values on the bar graphs correspond to: n embryos that developed to blastocyst/total n embryos in the group. **i,** Average ICM:TE ratio in control versus miR-221-3p mimic-treated blastocysts, where each data point represents the ICM:TE ratio of one embryo. **j,** Representative immunofluorescence and brightfield (BF) images of control (n=17) and miR-221-3p mimic-treated (n=9) blastocysts. Treatment/staining/imaging was performed in 4 separate batches. Hoechst = nuclear stain. Comp = composite image. All statistical comparisons were conducted using either a One-Way ANOVA or non-parametric (Kruskal-Wallis) test where appropriate. Values are reported as means ± standard error. *** = *p* < 0.001.

We then aimed to identify key miRNAs involved in driving lineage specification or maintenance by applying trajectory and pseudotime analyses to reconstruct dynamic processes and model continuous cellular changes. A clear bifurcation was identified beginning at the 8-cell and morula stage, creating one trajectory for the ICM and one for the TE (Fig. 4b-f and Supplementary Data 4). miRNAs that were highly expressed prior to the establishment of lineage before decreasing with developmental progression included specific members of the let-7 family as well as miR-100-5p, miR-98-5p and miR-465b-5p (Fig. 4g, Extended Data Fig. 4a and Supplementary Data 4). Development of the ICM trajectory was marked both by the upregulation of several key miRNAs already expressed at lower levels in earlier stages from the *Dlk1-Dio3* locus (e.g., miR-127-3p, miR-370-3p, miR-381-3p) and by the emergence of new miRNAs not previously detected (miR-302 family) (Fig. 4g, Extended Data Fig. 4a). When examining the dynamics in the TE trajectory, we observed the upregulation of several miRNAs coinciding with TE formation. However, this increase was not entirely unique to the TE. Instead, it reflected a broader blastocyst-wide rise in expression, as these miRNAs were also upregulated in ICM cells, albeit to a lesser extent (e.g., miR-744-5p, miR-429-3p, miR-125a-5p, miR-467) (Fig. 4g, Extended Data Fig. 4a and Supplementary Data 4). In addition, a group of miRNAs were identified that underwent a very brief surge specifically in the ‘emergent TE’ followed by a significant decrease in the early to middle TE (e.g., miR-221-3p, miR-223-3p, miR-203-3p, mir-205-5p, miR-27a-3p) (Fig. 4g, Extended Data Fig. 4a and Supplementary Data 4). Overall, our data suggests there is a strong miRNA signature exclusively associated with the ICM lineage trajectory, while fewer miRNAs distinguish the TE trajectory. Instead, a “surge” of miRNAs early in TE specification (in late morulas and very early blastocysts) may contribute to mouse embryonic cells adopting a TE fate.

We next investigated how the transient surge of emergent TE-associated miRNAs might influence TE specification and overall preimplantation development. Because miR-221-3p was among the highest expressed at this stage (Fig. 4g) and has been previously characterized as an anti-stemness miRNA^45,46^, it was selected for functional testing. We introduced a miR-221-3p mirVana miRNA mimic (Invitrogen™, Thermo Fisher Scientific) or a scrambled negative control mimic into the culture media of 2-cell stage embryos at final concentrations of 5 µM or 10 µM. Treatment with 10 µM, but not 5 µM of miR-221-3p resulted in a significant decrease in the rate at which embryos developed to the blastocyst stage by E3.5 compared to controls (Fig. 4h). Mimic-treated embryos stalled in development between the morula and mid-blastocyst stage, with embryos showing visual signs of poor quality (lack of blastocoel, cell fragmentation, embryo degradation), while control embryos developed to blastocysts and began hatching (Extended Data Fig. 4b). Further, examining the lineage proportions in E3.5 embryos (Sox2 (ICM marker) and Cdx2 (TE marker)) we observed that the 10 µM dose skewed the number of cells belonging to each lineage, causing a significant increase in the number of TE cells and a decrease in ICM cells (Fig. 4i, Extended Data Fig. 4c). Notably, this result was not due to a change in the total cell number of embryos at this stage, which remained consistent between treatment and control (Extended Data Fig. 4d).

Quantification of the average nuclear fluorescence intensity of Sox2 protein in ICM cells demonstrated that miR-221-3p mimic also decreased its protein expression in a dose-dependent manner compared to controls (Extended Data Fig. 4e). This is consistent with previous findings in mESCs, where transfection of miR-221-3p mimic induced spontaneous differentiation by targeting of the 3′ untranslated regions of mRNA transcripts of the major pluripotency factors Oct4, Nanog, and Sox2^45^. Unexpected however was a dose-dependent decrease in the expression of Cdx2 in TE cells (Extended Data Fig. 4f), which was not a predicted target. This decrease in Cdx2 expression may be a consequence of the targeting of Sox2, as Sox2 siRNA-treated embryos fail to form TE and have downregulated expression of TE markers, including Cdx2, Tead4, Yap and Eomes^60^. Overall, these results suggest that precocious expression of miR-221-3p disrupts preimplantation development and blastocyst formation by skewing lineage allocation, likely through suppression of Sox2 and other downstream effects on both ICM maintenance and TE identity. Further detailed studies are required to determine if Sox2 is in fact a direct target of miR-221-3p in the mouse embryo, and the exact mechanisms through which other miRNAs of the emergent TE “surge” may act to promote TE specification.

### miRNA target analysis and potential influence on lineage specification

Canonically, mature miRNAs achieve post-transcriptional gene expression regulation by associating with Argonaute proteins to form the miRNA-induced silencing complex (miRISC), which then binds complementary sites in the 3′-UTRs of target mRNAs via Watson–Crick base pairing, triggering translational repression or mRNA degradation^13,61^. Given that miRNAs can share similar expression patterns (Fig. 4e-g and Extended Data Fig. 4a), that one miRNA may have numerous targets, and that separate miRNAs can share the same target and act in concert, we classified miRNAs based on their coexpression patterns using single-cell weighted gene coexpression analysis (scWGCNA) applied to the full-cell Small-Seq data^62,63^. This approach allowed us to investigate the potential functional roles of co-expressed miRNA clusters during mouse embryonic development. In total, we identified 10 distinct miRNA modules, whose robustness was confirmed by module preservation analysis (Fig. 5a,b and Extended Data Fig. 5b) with four differential module eigengenes (DMEs) between ICM and TE. Notably, 84.1% (142 out of 169) of trajectory-related miRNA were included in the different modules (Fig. 5c and Supplementary Data 5). In addition, 134 extra differentially expressed miRNA were also included, indicating that scWGCNA effectively represents the overall dynamics of miRNA expression (Supplementary Data 5).

**Fig. 5.**
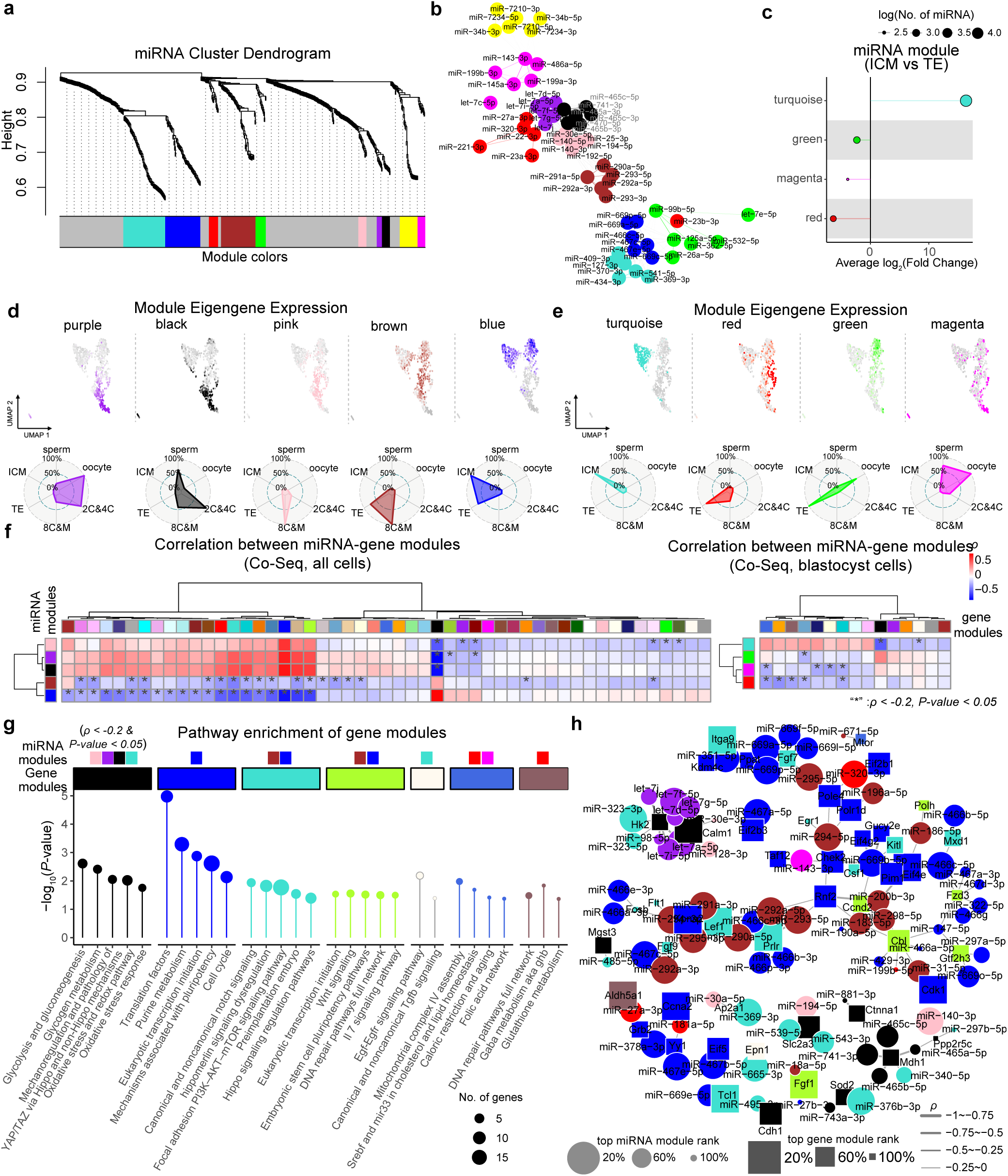
Weighted gene co-expression network analysis (WGCNA) of miRNAs in the mouse preimplantation embryo. **a,** Cluster dendrogram detailing identification of miRNA modules identified in full cells of the main Small-Seq dataset. Module colors are indicated at the bottom, with grey representing genes not assigned to any co-expression module. **b,** Network visualization of hub miRNA within the miRNA co-expression module. Node size is proportional to intramodular connectivity (kME), and line thickness represents the strength of co-expression between miRNAs. Highly connected hub miRNAs are highlighted. **c,** Significant differential module eigengenes (DMEs, P-value < 0.05) between the ICM and TE. The x-axis shows the average log₂ fold change (ICM vs. TE) for each module. Dot color corresponds to the assigned module, and dot size reflects the log-transformed number of miRNAs within the module. UMAPs and radar plots showing **d,** non-lineage specific and **e,** lineage-specific miRNA module expression across all cells based on Small-Seq data. **f,** Spearman correlation (ρ) between miRNA modules and gene modules from the same cells in Co-Seq data, where significant negative correlations (blue) are marked with a “*” (ρ< -0.2 and p-value < 0.05). **g,** Pathway enrichment analysis of gene modules that were significantly negatively correlated with the indicated miRNA modules. Dot color corresponds to the assigned gene module, and dot size signifies the number of genes within the module related to that pathway. Color of miRNA modules with significant negative correlation were labeled above. **h,** The network plot shows miRNA–target gene relationships from significantly correlated modules. Only miRNA–target gene pairs with significant negative correlations (p-value < 0.05) and genes belonging to enriched pathways were included. For miRNA–gene module pairs with more than 15 connections, only the top 15 pairs (ranked by module eigengene-based connectivity(kME)) were retained. Circles represent miRNAs and are colored according to their module; squares represent genes and are colored according to their respective modules. The size of each circle or square reflects the kME rank within its module, and line thickness indicates the Spearman correlation (ρ).

Among the identified modules, the yellow module—with hub miRNAs miR-34b-3p and miR-7210, were predominantly expressed in sperm cells (Fig. 5b and Extended Data Fig. 5a). The purple module, enriched for the let-7 family as hub miRNAs, showed high expression in oocytes as well as at the 2-cell and 4-cell stages (Fig. 5b,d). The black module, with miR-465 as its hub, was enriched at the 2-cell and 4-cell stages and also expressed at relatively high levels in sperm. The pink and brown modules were mainly expressed in 8-cell and morula cells. In addition, the blue module, comprising miR-669 and miR-467 from the C2MC cluster, was highly expressed across both blastocyst lineages (Fig. 5b,d), consistent with our previous analysis above. Notably, among the DMEs distinguishing ICM from TE, the turquoise module—containing C12MC members such as miR-370 and miR-409—was preferentially expressed in ICM (Fig. 5b–d). In contrast, the red, green, and magenta modules were predominantly expressed in TE (Fig. 5c-d). To utilize the Co-Seq data for a systematic investigation of gene–miRNA expression relationships, we projected the identified miRNA modules onto the miRNA expression profiles from Co-Seq (Extended Data Fig. 5b and Methods). Excluding the yellow module (as sperm was not sequenced using Co-seq), all other modules exhibited preservation between full Small-Seq and the Small-Seq portion of Co-Seq (Extended Data Fig. 5b), further supporting the robustness of these co-expression miRNA modules and the high quality of our Co-Seq data. Similarly, co-expressed genes were categorized into 57 modules (Extended Data Fig. 5c), of which 16 represented DMEs distinguishing ICM from TE (Extended Data Fig. 5d).

Spearman correlations were then calculated between all the miRNA modules and gene modules (Fig. 5f and Supplementary Data 6). The top negatively correlated gene–miRNA module pairs include black-gene/black-miRNA and blue-gene/blue-miRNA (Extended Data Fig. 5e and Supplementary Data 6). miRNA target enrichment within each module pair further supported that these negative correlations were likely caused by direct targeting rather than secondary or downstream regulation (Extended Data Fig. 5f). The negative correlation between black-gene and black-miRNA modules was mainly attributable to miR-465, while the blue-gene/blue-miRNA pair was driven by the miR-466/467 family (Extended Data Fig. 5f). In addition, we found that the blue miRNA module, which includes C2MC miRNAs as hub regulators, showed opposite expression to the black miRNA module and exerted broad negative influences on multiple gene expression modules, suggesting its wide regulatory roles during very early zygote and morula development. Further, the ICM-enriched turquoise miRNA module was predicted to regulate the TE-enriched black gene module, (Fig. 5f,g and Extended Data Fig. 5d), primarily through the activity of miR-495 (C2MC), miR-539, and miR-377 (C12MC) (Extended Data Fig. 5f), supporting our notion that these miRNAs may be important for pluripotency and embryo development.

Using curated databases (miRDB^64^ and TargetScan^65^), we inferred genome-wide targets for each miRNA module. We then focused on predicted targets that were negatively correlated with their corresponding miRNA modules and performed pathway enrichment analysis to uncover potential functional roles of these miRNAs. Notably, miRNA modules corresponding temporally with blastocyst formation (brown, blue, pink, turquoise) were negatively correlated with and predicted to target genes from gene modules (black and turquoise gene modules) related to the Hippo signaling pathway (Fig. 5g). As differential activation of Hippo/YAP signaling is known to drive ICM–TE specification in the mouse embryo^1^, this could suggest that miRNAs contribute to this process by fine-tuning the expression of key components, thereby promoting the acquisition of distinct cell identities. Another interesting observation is the presence of genes related to glycolysis/gluconeogenesis pathways among the black gene module, which are negatively correlated with the pink, purple, black and turquoise miRNA modules (Fig. 5g). Differences in glycolytic activity and the expression of glycolytic genes are known to occur upon blastocyst formation^66^. In the TE, glycolysis promotes lineage specification by driving the nuclear localization of Yap and the translation of *Tfap2c*, whereas it does not appear to play a role in the ICM^67^. Consistent with this, we found that miRNAs from the turquoise ICM module (e.g., miR-323-3p, miR-323-5p) are predicted to target and suppress *Hk2* (Fig. 5h), a critical enzyme catalyzing glycolysis. These findings indicate that multiple miRNAs could modulate glycolytic pathways and Hippo/YAP signaling during mouse preimplantation development and may contribute to ICM–TE specification, but the roles of specific miRNAs and genes remain speculative and will require further functional studies to validate.

To validate our miRNA–gene module regulation analysis, we reanalyzed a previously published single-cell transcriptomic dataset from our lab, in which mouse embryos were cultured from the 2-cell stage through to the 32-cell blastocyst stage in the presence of a miR-381-3p mimic^14^. This dataset was originally generated to assess how miR-381-3p, an ICM-enriched miRNA in mouse and human^14^ (*Dlk1-Dio3* locus), may influence global gene expression in the blastocyst. Applying the gene modules identified in the current study to the re-analysis of this dataset, we observed a significant downregulation of transcripts specifically belonging to the black gene module in miR-381-3p-treated blastocysts compared to controls (Extended Data Fig. 5g). Because miR-381-3p belongs to the turquoise miRNA module, which is negatively correlated with and predicted to target the black gene module, these results support our target analysis and indicate that *in silico* predictions can guide future functional studies, while also demonstrating that miRNA mimics act specifically within embryos.

It is possible that stage- and lineage-specific co-expression of miRNAs may reflect regulation by shared transcription factors (TFs)^36,62,68^. To investigate this, we identified TFs with enriched binding sites in known regulatory regions of each miRNA module (Fisher’s exact test: odds ratio [OR] > 1, p < 0.05; Extended Data Fig. 6 and Supplementary Data 5) based on TransmiR v3.0^68^, a database containing literature-curated and ChIP-seq–derived TF–miRNA regulatory interactions. TFs potentially regulating the blue miRNA module (enriched in all blastocyst cells) can be divided into two types. The first type is related to stemness, including *Nanog*, *Zfp281*, *Rest*, and *Yy1* (Extended Data Fig. 6a,b and Supplementary Data 5). The second type consists of epigenetic or complex-associated TFs, such as components of the Polycomb Repressive Complex (Extended Data Fig. 6a,b). In contrast, TFs likely to regulate the green miRNA module (specifically enriched in TE) included *Cdx2* and *Gata6,* which are known to contribute to TE and PE specification, respectively (Extended Data Fig. 6a,c). These findings suggest that, beyond their established roles in lineage specification^1^, these TFs may further reinforce ICM or TE identity by activating key lineage-associated miRNAs and potentially create a regulatory feedback loop. Also of note, *Yy1* was predicted to regulate the turquoise (enriched in ICM) miRNA module (Extended Data Fig. 6a,d). In recent studies, shaping of nucleosome organization and enhancer accessibility by *Yy1* has been shown to be essential for early mouse embryo development^69,70^ and for ESC differentiation into the three germ layers^71,72^. It is therefore plausible that such chromatin regulatory activity may facilitate the expression of miRNAs critical for ICM and TE differentiation potential; however, further work will be required to determine whether these mechanisms operate in the preimplantation embryo to reinforce cell fate decisions.

### Other sncRNAs in mouse preimplantation development

Next, we plotted and analyzed the single-cell expression patterns of the other sncRNA subtypes that are detected with Small-Seq (snoRNA, tRNA, piRNA, snRNA, rRNA) in mouse germ cells and preimplantation embryos, to determine if any apart from miRNAs display developmental stage- or lineage-specific expression signatures. We found that sperm cells formed distinct clusters based on the expression of the majority of sncRNA subtypes, suggesting they have very distinguished sncRNA expression profiles from oocytes and preimplantation embryos (Extended Data Fig. 7a,b). In addition, oocytes, 2-cell, and 4-cell stage embryos seem to have a different piRNA profile than blastocyst cells (Extended Data Fig. 7a,b). However, we only observed distinct clustering based on both developmental progression and blastocyst cell type (ICM and TE clusters) for snoRNA expression (Fig. 3c and Extended Data Fig. 7a-c), similar to what we have previously shown in human preimplantation embryos^14^. In particular, a very strong snoRNA signature was found in ICM cells, segregating them from TE cells (Extended Data Fig. 7c and Supplementary Data 2). Notably, many of these snoRNAs arise from the *Dlk1-Dio3* locus (C12MC), similar to ICM-specific miRNAs (Fig. 3c). Although the potential role of snoRNAs in pluripotency/differentiation has not been thoroughly investigated, one study has shown that some H/ACA snoRNAs (base pair with targets such as rRNAs, snRNAs and mRNAs leading to their pseudourdylation) are differentially expressed during mESC differentiation^73^. In addition, snoRNAs are known to regulate spliceosome function, and pluripotent cells have globally elevated spliceosome activity compared with differentiated cells^74,75^. Overall, it is plausible that snoRNAs may contribute to the maintenance of pluripotency in the ICM by a wide variety of mechanisms, but further studies are required to determine exactly how specific snoRNAs may contribute to lineage specification in the embryo.

### sncRNA inheritance from sperm and oocyte to early embryo

Investigation of sncRNA inheritance from gametes to early embryos can provide insights into parental contributions to epigenetic regulation, embryonic lineage specification, and potential links to developmental disorders^76–78^. As such, we sought to identify which sncRNAs are expressed amongst sperm, oocytes, and 2-cell embryos from our mouse Small-Seq dataset, to see which may be maintained post-fertilization. This type of investigation has previously been performed on whole mouse embryos by real-time PCR-based or sequencing methods to identify sncRNAs, including miRNAs, which are expressed and may be inherited^32,78–83^, however, we reasoned that single-cell profiling of all the sncRNAs would provide deeper insight even in regards to miRNAs, as Small-Seq simultaneously captures all sncRNAs and reduces amplification-related technical noise through the use of unique molecular identifiers (UMIs).

Based on quantification of UMIs, we detected a total of 584 miRNAs in the oocyte (Extended Data Fig. 8a and Supplementary Data 7). Using stringent parameters to minimize the detection of sncRNAs with low expression frequency and generate false positives (see Methods), from these, we found 286 miRNAs co-expressed amongst oocytes, sperm and the 2-cell embryo, with an additional 10 miRNAs co-expressed exclusively between the 2-cell embryo and oocytes (e.g., miR-291) (Extended Data Fig. 8a-c and Supplementary Data 7). Among these is miR-99a-5p, which suggests that it is inherited (Supplementary Data 7) in-line with previous works^32^. Notably, among the oocyte-inherited miRNAs were also members of the miR-290-295 family (e.g., miR-291a, miR-294, miR-295) (Supplementary Data 7 and Extended Data Fig. 8d). It has previously been reported that miR-290-295 miRNAs undergo extensive 3’oligoadenylation at the 1- and 2-cell stages^32^, which may function to stabilize them during the large-scale miRNA degradation occurring between oocyte and zygote^79^. Together, these 296 (286+10) miRNAs likely represent germ cell inherited miRNAs since de novo production is believed to commence during the 2-cell stage, coinciding with the major zygotic genome activation^84^. Consistent with this, we only observe 2 miRNAs (e.g., miR-215-5p, miR-7b-5p), which are exclusively expressed in the 2-cell embryo (Supplementary Data 7 and Extended Data Fig. 8d). Our results are in line with previous observations which show that miRNA profiles of the oocyte and the 2-cell embryo are similar^32,79^ (Fig. 2c and Extended Data Fig. 7).

In sperm, miRNAs are believed to be of somatic or mixed origin, but nonetheless have been shown to have potential impacts on embryogenesis^32,76,85^. We detected a total of 454 miRNAs in sperm based on the initial quantification of UMIs (see Methods) (Extended Data Fig. 8a, and Supplementary Data 7). Applying similar stringent parameters, as with the oocyte, to minimize false positives when determining exclusive lineage expression, we identify 20 miRNAs expressed exclusively in sperm (Extended Data Fig. 8b,c and Supplementary Data 7). These include the previously reported miR-34c-3p, which has been shown to be highly expressed in mouse spermatids and human sperm^85–89^ and miR-7210, miR-7217, the miR-7232–7234 cluster, and the miR-7241/7242 cluster, whose functions remain to be studied (Supplementary Data 7). We also detected a well known sperm miRNA, miR-449a-5p, as co-expressed between the oocyte, sperm and 2-cell embryo, in-line with previous work demonstrating its inheritance from sperm to the 1-cell embryo^86^ (Supplementary Data 7). Given our stringent parameters, we detected 0 miRNAs exclusively shared between sperm and 2-cell embryos, which is likely an underrepresentation; however, as mentioned above, 286 miRNAs were shared amongst sperm, oocyte and 2-cell embryos, representing the payload from sperm; consistent with prior studies that have shown miRNAs inherited from sperm regulate proper maternal mRNA turnover during the zygote to 2-cell transition and the first cleavage division^77,87^. Further, previous work has shown that *Dgcr8*-deficient oocytes fertilized with sperm was sufficient to produce viable pups^90^, given the considerable overlap we observe between sperm and oocyte miRNAs, perhaps the paternal or maternal contribution is sufficient and has a redundant role in development to serve as a ‘fail safe’ mechanism. Importantly, our *in silico* analysis cannot determine with certainty whether the shared miRNAs identified amongst the oocyte, sperm and 2-cell embryo persisted during the 1- to 2-cell transition, representing bona fide inheritance, as functional experiments verifying their inheritance was not performed. Nonetheless, understanding the dynamics of miRNAs during fertilization and embryo formation is of interest, particularly in light of recent works in the fields of Developmental Origins of Health and Disease showing the impact of maternal and paternal lifestyle on embryo and offspring health.

tRNAs are known to be produced in both sperm and oocytes, which post-fertilization are utilized for *de novo* protein synthesis^32,78,91–93^. There is also growing evidence that within the early embryo, tRNA fragments regulate expression of transcripts driven by endogenous retroelements^94^. We identified 436 tRNAs and tRNA fragments shared amongst the sperm, oocytes and 2-cell embryo (Extended Data Fig. 8a-c and Supplementary Data 7). Indeed, amongst those which overlap with germ cells and the 2-cell embryo, included 15 tRNA belonging to tRNA-Gly-GCC family, which have been shown to regulate embryonic gene transcription by promoter binding, contribute to genomic stability via transposon silencing and potentially transmit parental environmental exposures to offspring for epigenetic inheritance^91,92,95–97^. Further, we identify 4 tRNAs, including tRNA-Leu-TAA-4-1specifically expressed in sperm (Extended Data Fig. 8b,e and Supplementary Data 7), as has previously been reported in human sperm^89^. Similar expression dynamics are observed for snRNAs and snoRNAs (Extended Data Fig. 8a-c and Supplementary Data 7). Finally, we observe zero tRNAs, snRNAs, snoRNA or rRNAs exclusively expressed at the 2-cell stage embryo, suggesting a lack of de novo production in the embryo for these sncRNA biotypes (Extended Data Fig. 8b,c). This is consistent with previous reports which have shown that rRNA biogenesis for instance begins at the 2-cell stage^98^. It is important to note that sperm RNA load is several fold lower than that detected in the oocyte or embryo. As such, it is possible that we have under-represented the number of sncRNAs expressed in our conservative analysis.

Finally, we examined the expression dynamics of piRNAs and consistent with previous reports, piRNAs are the most abundant sncRNA in sperm and are believed to be of true germ cell origin^76^. Indeed, we detect 42,276 piRNAs expressed in sperm and 14,203 piRNAs expressed in the oocyte (Extended Data Fig. 8a and Supplementary Data 7). Applying the same stringent parameters to minimize false positives when describing exclusive expression, among the differentially expressed piRNAs between sperm, oocytes, and 2-cell embryos, at least 27,734 piRNAs (52.0%) were found exclusively in sperm, whereas 155 piRNAs, were found to be oocyte specific and 1 piRNA (piR-mmu-5029435) was exclusively shared between the oocyte and sperm but not passed onto the 2-cell embryo (Extended Data Fig. 8b,c and Supplementary Data 7). In contrast, 1287 piRNAs are detected at the 2-cell stage overlapping with sperm and oocyte expression with an additional 1495 piRNAs shared specifically with the oocyte (Extended Data Fig. 8b,c and Supplementary Data 7); in-line with previous work suggesting inheritance of piRNAs from germ cells^99^. Further, we only detected 5 piRNAs exclusively expressed in the 2-cell embryos, suggesting de novo production (Extended Data Fig. 8b,c and Supplementary Data 7). The abundant piRNAs inherited from the oocyte to the early embryo could be an important mechanism given the loss of expression of piRNA biogenesis players very early in preimplantation development (Fig. 1e and Extended Data Fig. 8b). Among the piRNAs which were explicitly inherited from oocyte (1495 piRNAs), the largest proportion of piRNAs originate from a region on chromosome 10 (325 piRNAs), located downstream of the protein-coding gene D10Wsu102e (Extended Data Fig. 8f). These D10Wsu102e-related piRNAs have been reported to show decreased expression during aging and may possess potential trans-regulatory functions^100^. We also observe 17 piRNAs, also enriched downstream of D10Wsu102e, which appear to be exclusively expressed in the oocyte (Extended Data Fig. 8f). While our data provides detailed insights into specific sncRNAs expressed both within the germ cells and likely inherited in the embryo, their roles remain to be further investigated.

### Cross-species comparison between mouse and human sncRNA sequencing results

An expanding body of research has revealed that despite superficial morphological similarities, mouse and human preimplantation embryos exhibit many differences in the timing, gene expression patterns, and signaling pathways underlying preimplantation lineage specification^1^. As such, we next sought to determine to what degree the commonly used mouse model recapitulates the human in terms of sncRNA expression dynamics by comparing the dataset in this study with an identical generated dataset from human preimplantation embryos from our group^14^. We identified 153 miRNA families (and 313 individual miRNAs, 19.6% of total miRNAs) that are expressed in both human and mouse preimplantation development (Fig. 6a). Among the miRNAs that were conserved in both species, 51.6% (157/313) were associated with developmental stage and lineage. For example, miRNAs from the *Dlk1– Dio3* locus (C12MC in mouse, C14MC in human) and the miR-302/367 family were specifically upregulated in the ICM of blastocysts in both species (Fig. 6b,c), although stratifying the lineages further by developmental progression revealed subtle differences in their expression dynamics (Fig. 6b,c). In the mouse, ICM miRNAs like miR-369-3 and miR−377−5p are already upregulated by the early blastocyst stage (E3.5) and continue to rise in expression by the mid blastocyst stage (E4.0) (Fig. 6c). In contrast, these same miRNAs remain at much lower levels in the human early blastocyst (E5) and only show a sharp increase at the mid- to late-blastocyst stage (E6/7) (Fig. 6c). The molecular basis for these species-specific differences in the timing of ICM miRNA expression remains to be determined. Notably, there were overall less miRNAs that were conserved between the mouse and human TE program (Fig. 6b,c), likely due in part to the overall lower amount of conserved TE-specific miRNAs in both species. Among the few that were consistent were miR-200c-3p and miR-24-3p (Fig. 6c), which have both been identified as differentiation-inducing miRNAs, targeting pluripotency factors like *Sox2, Oct4, Foxo1, c-Myc* and *Smads*^101–103^. In addition, miR-200c-3p has been shown to regulate the Hippo signaling pathway by targeting *Lats1/2*^104^, which mechanistically fits in the TE as this would promote the nuclear localization of Yap ultimately supporting TE cell fate.

**Fig. 6.**
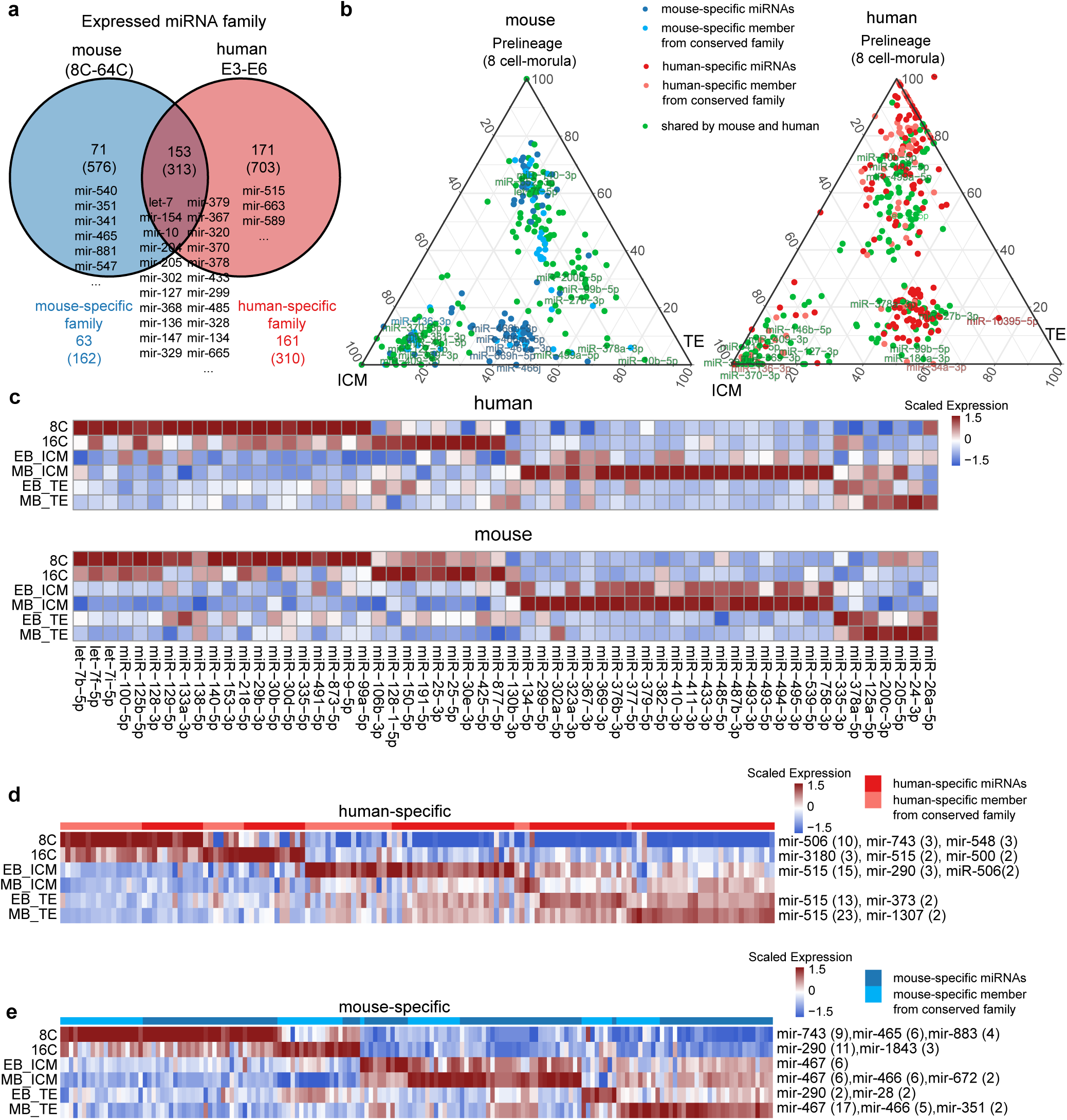
Cross-species comparison of miRNA expression dynamics in the mouse and the human preimplantation embryo. **a,** Venn diagram showing the overlap of expressed miRNA families between mouse and human embryos. The number of individual miRNAs is indicated in parentheses. For miRNA families and individual miRNAs from those families detected only in mouse or human, the number specific to each species, based on annotation, is highlighted. **b,** Ternary plot showing the average expression of differentially expressed miRNAs in prelineage, ICM, and TE. Based on annotation, miRNAs are colored according to their presence in the human and mouse genomes. **c,** Heatmap showing miRNA expression patterns reveals similar trends between mouse and human. **d-e**, Heatmaps displaying the average expression of lineage-related miRNAs in human (**d**) and mouse (**e**). Species-specific miRNAs, either representing distinct members of conserved miRNA families or being uniquely species-specific, are indicated by the column colors at the top. On the right, the top three miRNA families, with at least two differentially expressed (DE) members, that are highly expressed within the same lineages and developmental stages. Numbers in parentheses denote the number of miRNA members within each family.

Although many key miRNA families displayed conserved expression, we also detected species-specific miRNAs expressed exclusively in either mouse or human. In total, 71 families (576 individual miRNAs) were expressed exclusively in mouse, whereas 171 families (703 individual miRNAs) were expressed exclusively in human (Fig. 6a). Of these, 33.3% (192/576) of mouse-specific and 22.3% (157/703) of human-specific miRNAs were previously linked to specific developmental stages or lineages^14^. We focused on these for further analysis, with the goal of pinpointing species-specific candidates that may function during preimplantation development and potentially reflect evolutionarily conserved or divergent regulatory mechanisms underlying blastocyst formation. Unsurprisingly, we identified miRNAs from the human-specific C19MC cluster (e.g., miR-515) (Fig. 6d and Supplementary Data 8) in the human TE but not the mouse. In addition, the rodent-specific miRNAs from the *Smfbt2* gene locus (C2MC; e.g., miR-466, miR-467, miR-669), which were highly expressed in the mouse ICM and TE, were not found in human (Fig. 6b,e and Supplementary Data 8). It is possible that C2MC represents a critical species difference, as we previously showed in our module and gene target enrichment analysis that these miRNAs are likely to have important functions during blastocyst development (Fig. 5b,d and Extended Data Fig. 4d). Another important distinction is the presence of the miR-322 found in mouse TE and shown to be critical for mouse TSC differentiation^50^, while its functional ortholog in human (miR-424) was not enriched in human TE (Fig. 5d,e and Supplementary Data 8). Overall, we present a comprehensive list of miRNAs which appear to be species-specific in early embryogenesis. These differences are important to consider when translating functional findings from mouse to human work, including human ESCs. Given the differential expression dynamics and lineage enrichment of miRNAs originating from both the C2MC and C12MC compared to the human counterparts C19MC and C14MC, we conclude that caution is warranted with the translatability of results obtained from functional studies during early mouse embryogenesis toward human application.

### Blastoid sncRNA comparison with the human blastocyst

Stem cell-derived structures modelling the human preimplantation blastocyst (Blastoids) have recently been developed^16–22^, providing a method to better understand early human development while avoiding some of the ethical constraints and limitations of human blastocyst research. Moreover, blastoids may more faithfully recapitulate aspects of the human blastocyst than the commonly used mouse model, with the additional benefit of reducing reliance on animal experimentation. While it has been shown that these blastoids contain cellular lineages highly reminiscent of their in vivo counterparts based on transcriptomic signatures^105^, how well these models recapitulate the TE and ICM of human embryos based on sncRNA expression has not been extensively studied. As such, we collected 199 single cells from 3 human blastoids resembling human blastocysts from the Theunissen lab^16^ and sequenced them using Small-Seq, comparing this to our previously published atlas of sncRNAs in human preimplantation development^14^.

Similar to our analysis of mouse embryo data, we first implemented Co-Seq on an additional 143 cells (from the same 3 blastoids plus 4 additional blastoids) and identified EPI-like cell (ELC) and TE-like cell (TLC) populations based on well-established lineage marker gene expression from the mRNA portion (Fig. 7a, b). Potential post-implantation-lineage-like cell contamination was further ruled out by examining additional lineage-specific markers (Extended Data Fig. 9a) and projecting onto a comprehensive embryonic reference using our Early Embryogenesis Prediction tool, proving similarities between blastoids and human blastocysts based on mRNA expression^105^ (Fig. 7c). Notably, cells with a PE transcriptomic signature were not detected in blastoids (Extended Data Fig. 9a and Fig. 7c), consistent with previous reports indicating that very few PE-like cells are generated in this model^16^. As in previous sections, after defining the cell identities based on gene expression, we further interrogated the miRNA expression of different lineage-like cells in the sncRNA portion of Co-Seq and integration with full Small-Seq cells allowed for annotation of the previously unidentified miRNA profiles (Extended Data Fig.9b and Fig. 7d).

**Fig. 7.**
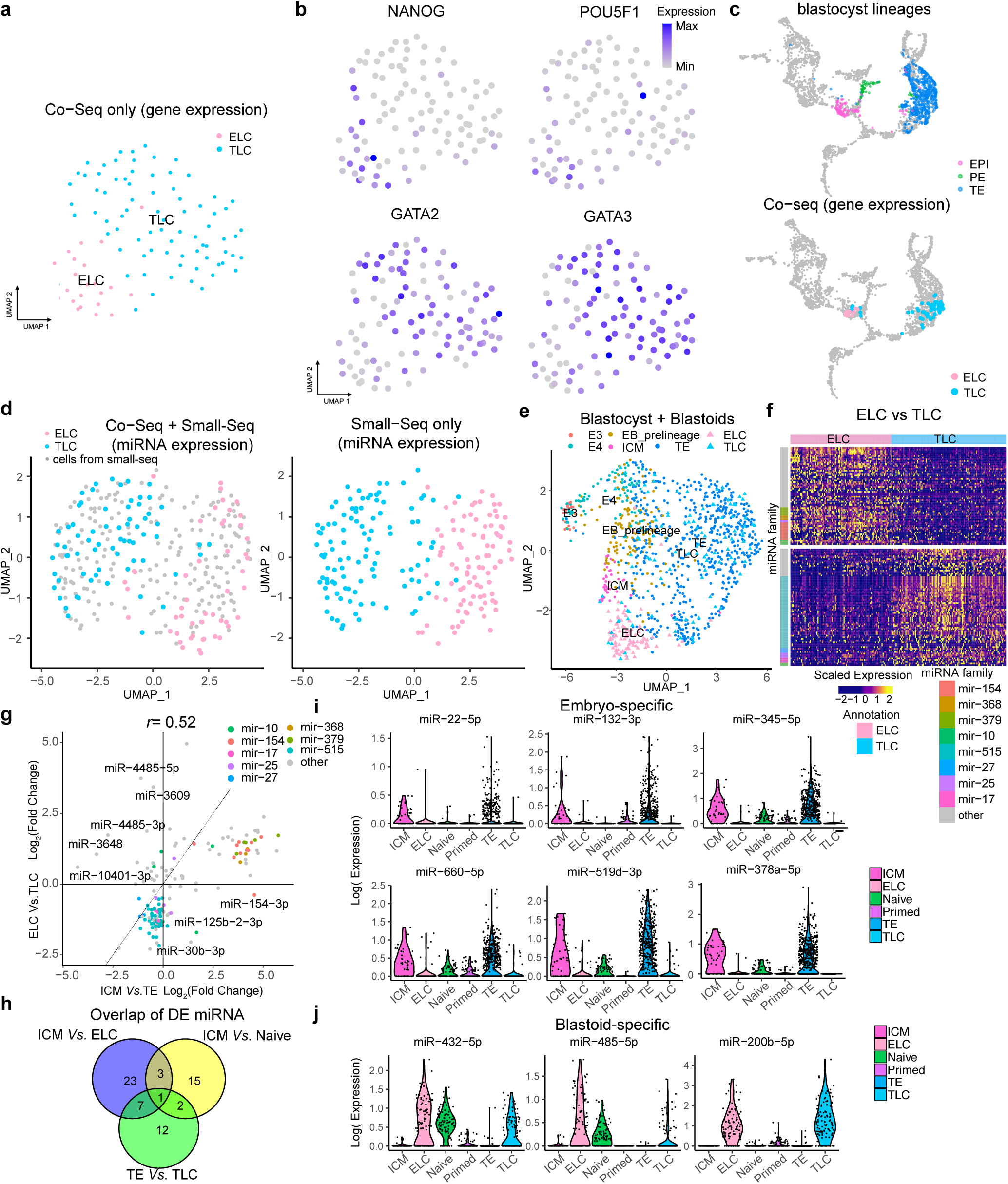
Lineage annotation and validation of human blastoid cells using gene and miRNA expression profiles. **a**, UMAP showing lineage annotation of human blastoids (mRNA portion from Co-Seq data) based on known transcriptome lineage markers. **b**, Expression of selected lineage-specific markers. ELC: *NANOG* and *POU5F1*; TLC: *GATA2* and *GATA3*. **c**, Projection of human blastoids (mRNA from Co-Seq data)) onto our human embryonic reference using the Early Embryogenesis Prediction tool^105^. **d**, UMAP showing integration of cells of known lineage identity (sequenced with Co-Seq) with unknown full cells of the human blastoids (sequenced with Small-Seq only). **e**, Integration of human blastoids miRNA expression obtained from Co-Seq (Small-Seq portion) full cell experiments. **f**, Heatmap displaying expression of differentially expressed (DE) miRNAs between ELC and TLC. miRNA families with at least three members that are highly expressed in ELC or TLC are indicated by the colors on the left. **g**, Scatter plot showing the log2(fold change) of DE miRNAs from the union set of ICM vs TE and ELC vs TLC. The x and y axis represents log2(fold change) of ICM vs TE and ELC vs TLC, respectively. Dot colors indicate miRNA families with at least three DE members. miRNAs with the most inconsistent expression patterns between ICM vs TE and ELC vs TLC are highlighted. **h**, Venn diagram showing the overlap of DE miRNAs among ICM vs ELC, ICM vs Naïve, and TE vs TLC comparisons. **i**-**j,** Violin plot showing the log-normalized expression of selected miRNAs only expressed in embryo (**i**) or blastoids (**j**) across ICM, ELC, Naive, Primed, TE and TLC. ELC: Epiblast-like cells. TLC: Trophectoderm-like stem cells

To assess how well blastoids recapitulate human blastocysts based on miRNA expression, we integrated our full-cell Small-Seq blastoid data with preimplantation human embryonic reference data^14^. As expected, human blastoids formed two separate clusters, integrating closely with human blastocyst cells based on their miRNA expression profiles. Indeed, blastoid TLCs clustered tightly with human TE, whereas blastoid ELCs exhibited miRNA profiles that clustered near, but not directly with, the human ICM (Fig. 7e). This divergence may be reflective of the lower number of miRNAs expressed in the blastoid ELCs when compared to the human ICM, despite the 63.7% overlap in miRNAs or the retainment of naive hESCs related miRNAs which are observed in the ELC (Fig. 7j and Extended Data Fig. 9c).

To further investigate the similarities between blastoids and human embryos, we examined DE miRNAs between ELCs and TLCs, and compared them with the DE miRNAs between the ICM and TE (Fig.7f and Supplementary Data 9). We found that miRNA families such as C14MC members (*Dlk1–Dio3* locus; e.g., miR-154, miR-368, miR-379) and C19MC members (e.g., miR-515) were preferentially enriched in the ELC and TLC, respectively (Fig. 7f), suggesting that these miRNA profiles may contribute to cell fate in a manner similar to the human embryo. A more detailed analysis of the log2 fold changes of DE miRNAs between ICM/TE and ELC/TLC showed considerable consistency (Spearman correlation r = 0.51) between the two datasets, with more pronounced differences in ICM-enriched miRNAs and TLC enriched miRNAs (Fig. 7g). Of particular interest, we noted enriched expression of miR-4485 in TLCs. MiR-4485 has been reported to regulate cell cycle progression and apoptosis by modulating mitochondrial function, leading to G2–M phase arrest and activation of caspase-mediated cell death^106,107^. This is in-line with recent work demonstrating the necessity of p53 and apoptosis in the differentiation^108^, though further work is required to confirm the interplay of miRNAs in this process.

We hypothesized that differences in individual miRNA expression between blastocysts and blastoids may reflect intrinsic stem cell-derived features that persist in blastoids. To investigate this, we compared the expression of blastocyst-specific and blastoid-specific miRNAs to a previously published^23^ Small-Seq dataset profiling the sncRNAs of naive and primed hESCs^23^ (Fig. 7i,j and Extended Data Fig. 10). Indeed, whereas blastocyst-specific miRNAs (e.g., miR-22-5p, miR-132-3p, miR-378a-5p) were low in naive and primed hESCs, some blastoid-specific miRNAs (e.g., miR-432-5p, miR-485-5p) were also characteristic of naive hESCs (Fig. 7i,j and Extended Data Fig. 10), suggesting that blastoid cells may retain certain miRNAs from their stem cell culture origins. Notably, blastocyst-specific miR-378a-5p has been implicated in trophoblast survival, migration, and invasion^110^, and its complementary strand, miR-378a-3p, has been reported to influence hatching in bovine blastocysts^109^. This may reflect a key difference between the blastoid model and the human embryo, particularly regarding TE behavior and implantation. Although beyond the scope of this study, future work could examine whether increasing these blastocyst-specific miRNAs in blastoids improves the functional fidelity of the model.

Finally, examining the other sncRNA subtypes, we found that similar to the human, snoRNAs segregate into two clusters based on cell lineage (Extended Data Fig. 9d). Indeed, 186 individual snoRNAs were differentially expressed between the blastoid ELCs and TLCs, including those from the *Dlk1-Dio3* locus (C14MC) enriched in ELCs (Extended Data Fig. 9e), as we have previously found in human blastocysts^14^. Although the function of snoRNAs originating from chromosome 14 in the human ICM and blastoid ELCs is not yet fully understood, the similarity in their expression patterns highlights blastoids as a promising model for uncovering the roles in governing pluripotency and the ICM and demonstrates similarities more closely to the human blastocyst than the mouse embryo in this context. In addition, we found that other sncRNA subtypes (tRNA and piRNA) also showed lineage-specific clustering, which we have not previously observed in human embryos^14^. The basis for this difference remains unclear, but we speculate that, as with the blastoid-specific miRNAs identified above, these signatures may reflect residual features carried over from their naïve stem cell origins. Nonetheless, these data can be leveraged to further optimize the semblance of blastoids to human blastocysts.

## Discussion

This study provides the first temporal single-cell atlas of sncRNA expression spanning mouse germ cells (oocytes and sperm) through preimplantation development (E1.5–E4.0). We demonstrate that among the major sncRNA subtypes, only miRNAs and snoRNAs exhibited stage- and lineage-specific expression patterns, distinguishing inner cell mass (ICM) from trophectoderm (TE) at the onset of the first cell fate decision. In contrast, piRNA expression underwent a broad shift at the blastocyst stage, which was not lineage-specific and, at least in part, likely reflected the overall decline in piRNA abundance that we and others^27,32^ have observed. As miRNAs are the most widely studied ncRNA in the context of development, our study delved deeper into examining their expression dynamics.

By the 32-cell stage, the ICM acquires a distinct miRNA profile enriched with miR-290-295, miR-17-92, miR-182-183 and *Dlk1-Dio3* (C12MC) miRNAs, all of which are hallmarks of naïve mESCs^40,58^. Maturation of the ICM into the EPI (64-cell stage) is also marked by a shift in miRNA profile, such as an increase in the miR-302 family, known to be more characteristic of primed mEpiSC^40,58^. All of these families function to sustain pluripotency through a wide variety of mechanisms in ESCs, which have been reviewed extensively^111^. For example, miRNAs from the *Dlk1-Dio3* locus repress genes that drive differentiation into the three germ layers (e.g., *Gata4*, *Foxa2*, *Sox17, Nestin*), and supplementation with members such as miR-541-5p, miR-410-3p, and miR-381-3p enhances the viability of ground-state ESCs^54^. Likewise, the miR-290-295 family regulates the cell cycle (G1/S transition via targets *Cdkn1a, Rb2* and *Lats2),* glycolysis (via target *Mbd2*), alternative splicing (via target *Mbnl1/2*), and the bivalent state of developmental genes, ultimately supporting maintenance of pluripotency^111^. Although functionally important in mESCs, *miR-290–295* knockout mice can survive until birth, albeit with female infertility in adults^112^. This suggests that it could be the concerted action of all the miRNA families mentioned above, and potentially additional miRNAs with overlapping functions (i.e., our identified miRNA modules), that is critical for robust ICM establishment and early embryonic development. Further, target and pathway enrichment analysis performed in this study identified additional potential functions of ICM miRNAs, including regulation of glycolysis/glucogenesis, oxidative stress and Hippo signaling. While we demonstrated that the ICM miRNA miR-381-3p (C12MC) can indeed reduce the expression of some predicted target genes in mouse blastocysts, further functional studies will be required to validate the other regulatory roles we have suggested.

In addition to our cross-species comparison, this study also provides the first single-cell characterization of sncRNA expression dynamics in the stem-cell based model of the human preimplantation blastocyst, the blastoid^16^, and compares it with human blastocysts. We found that the blastoids we profiled were largely similar to the E6 human embryos based on miRNA expression in the EPI-like and TE-like lineages, apart from a few miRNAs that were embryo-specific, and a few miRNAs that were blastoid-specific (possibly resulting from maintenance of miRNAs from their stem cell origins). Our results provide an additional molecular layer for benchmarking this model and add confidence to their future use for studies regarding miRNA function, while highlighting differences that may be important to take into consideration. One of the most striking of these differences was the lineage-specific clustering of sncRNA subtypes other than miRNAs and snoRNAs, which is not observed in the human blastocyst. A recent study performed multi-omics analysis of both embryos undergoing lineage specification, as well as induced pluripotent stem cells (iPSCs) undergoing reprogramming and TSC fate^113^. They reported that embryos follow a ‘T-shaped’ differentiation trajectory—in which cells initially undergo similar transcriptional changes before diverging into ICM and TE at the morula/early blastocyst stage—whereas reprogrammed cells follow a ‘V-shaped’ trajectory, adopting mutually exclusive chromatin and transcriptional programs from the onset of reprogramming^113^. While the sncRNAs were not profiled, it is possible that since this “V” pattern was conserved across the RNA, DNA, chromatin and methylation levels, that sncRNA expression also follows suit. Overall, the nature of cells being induced to differentiate (as occurs in the blastoid) may be a less gradual process than occurs in the natural embryo. Since the cells of the blastoid are also induced to differentiate via different methods/mechanisms, this explanation remains largely speculative, and would require further studies to be confirmed in the blastoid.

Although the extensive research that has been conducted allows for considerable speculation regarding the expression and functional roles of miRNAs, much less is known about snoRNAs, which have not been as thoroughly studied. Our finding of lineage-specific snoRNA expression in the mouse, human and blastoid—most notably the upregulation of Dlk1–Dio3 snoRNAs in the ICM—suggests that they may contribute to lineage acquisition, although their precise mechanisms remain unclear. The most well-known function of snoRNAs is to guide chemical modifications of other RNAs, primarily rRNAs and small nuclear RNAs snRNAs. However, recent studies have identified novel mechanisms for snoRNAs function, such as stabilizing target mRNAs^114^ and modulating mRNA 3’end processing regulating the expression of certain subsets of miRNAs^115^. In addition, snoRNAs are processed into smaller specifically stabilized fragments known as snoRNA-derived RNAs (sdRNAs), which are often conserved across species^115^. sdRNA production is widespread, with more than half of all snoRNAs shown to produce smaller fragments^115^. Notably, a predicted possible function for sdRNAs is as a novel source of miRNAs or piRNAs^115,116^.

Here we demonstrate dynamics of all sncRNAs in the mouse oocytes, sperm and throughout preimplantation embryogenesis. Further, we reveal similarities and highlight some important differences with the human embryo. Finally, we provide an additional benchmarking molecular layer for the blastoid and reveal close overlap with miRNAs, including C14MC and C19MC, and snoRNAs, providing further evidence that blastoids represent a powerful model and provide a useful tool, toward better understanding human embryogenesis. Collectively, this in-depth analysis advances our understanding of early embryonic development and paves the way for future studies aimed at targeted therapies in reproductive health and mechanistic studies to understand the interplay of epigenetics and genetics in blastocyst formation and pluripotency. Additionally, this sncRNA atlas can serve as a valuable reference for benchmarking stem cell–derived embryo models against their in vivo counterparts.

## Methods

### Ethics Statement

All animal experiments were approved by the Centre de Recherche du Centre Hospitalier de l’Université de Montréal (CRCHUM) Comité Institutionnel de Protection des Animaux du CHUM (CIPA) under the protocol number IP22026SPs. Blastoid experiments in the Theunissen lab were approved by the Embryonic Stem Cell Research Oversight (ESCRO) Committee at Washington University School of Medicine under protocol number 19-004.

### Collection of mouse preimplantation embryos, oocytes and sperm

MII oocytes and preimplantation embryos were collected from 6- to 8-week-old C57BL/6 female mice (Charles River Laboratories, Raleigh, NC, USA). The mice were delivered at 3 to 4 weeks of age, and animals were maintained in individually ventilated cages (up to 5 animals/cage) at 22±2°C and 40-60% humidity in 12 h light/dark cycle with lights on from 6:30am-6:30pm and ad libitum access to food and water. To induce ovulation, females were injected with 5 IU of pregnant mare serum gonadotropin (PMSG) followed by 5 IU human chorionic gonadotropin (hCG) 48 hours later^36,117,118^. To collect meiosis II (MII) oocytes, females were sacrificed using isoflurane anesthetic followed by cervical dislocation 12 to 14 hours following the hCG injection, and uteri were dissected and flushed. To collect 2-cell preimplantation embryos, female mice were placed with C57BL/6 males for mating immediately following the hCG injection and sacrificed as described above 45 to 48 hours later. In all cases, uteri were flushed with 37°C EmbryoMax M2 medium (Sigma-Aldrich, cat # MR-015-D) as we previously described^117,119^.Mouse sperm was collected following a previously described swim-out procedure^119^. Briefly, caudal epididymides of C57BL/6 males (obtained, housed, and euthanized as described above) were placed in Donner’s solution for sperm collection^119^, cut, and gently agitated at 37°C to allow sperm to swim out. The spermatozoa were then passed through a 40 μM micro-pore filter (Fisher Scientific, #22363547) to remove somatic cells. Finally, spermatozoa were transferred into freezing medium (Fujifilm, cat # 90128), and placed in a -80°C freezer for storage. To recover the sperm for downstream analysis, frozen samples were removed from the freezer and placed on ice. Once fully thawed, the sperm underwent three washes with PBS (by centrifuging at 2.6 g for 6 minutes to pellet the cells, then removing the supernatant). 10 µL was taken from the washed sperm cell suspension and loaded onto a hemocytometer to verify no somatic cell contamination and to count sperm heads (sperm counts: mouse 1 = 63.5*10^5^/mL, mouse 2 = 75.5*10^5^/mL, mouse 3 = 87.0*10^5^/mL).

### Embryo culture and treatment with mirVana miRNA mimics

Embryos that were to be sequenced with Small-Seq were were collected at the 2-cell stage and further cultured as we have previously described^117^, with each embryo isolated in its own drop of KSOM throughout, to rule out impacts on miRNA expression resulting from secretion and uptake from other embryos. Upon reaching the desired developmental stage (4-cell (54 to 56 hours post-hCG), 8-cell (68 to 70 hours post-hCG), 16-cell (78 to 80 hours post-hCG), 32-cell (92 to 94 hours post-hCG), and 64-cell (100-104 hours post-hCG)), embryos were removed from culture for single-cell dissociation and library preparation with Small-Seq.

To test the impact of increased expression of miR-221-3p during mouse preimplantation development, mirVana^TM^ custom miRNA mimics (Invitrogen^TM^ (ThermoFisher Scientific)) were utilized. MiR-221-3p mimic (Product ID# MC10337, Sequence: AGCUACAUUGUCUGCUGGGUUUC) or a scrambled negative control mimic (cat # 4464058) were added directly to KSOM media droplets (at final concentrations of 5 µM or 10 µM) containing group cultures of 2-cell embryos. We and others have demonstrated that treatment with mimics at the 2-cell stage results in passive uptake by embryos as early as the 4- and 8-cell stages^14,109,120^. Treated embryos were further until the 32-cell blastocyst stage.

### Human Blastoid Generation

Blastoid formation was performed as previously described by the Theunissen Lab^16^. Naïve H9 embryonic stem cells (ESCs) were cultured in 5i/L/A^121^ [1:1 DMEM/F12 (Gibco, 11320), 1:1 Neurobasal (Gibco, 21103), 1x N2 100X supplement (Gibco, 17502), 1x B27 50X supplement (Gibco, 17504), 1X GlutaMAX, 1X MEM NEAA (Gibco, 11140), 0.1 mM β-mercaptoethanol (Millipore Sigma, M3148), 1% penicillin-streptomycin (Gibco 15140-122), 50 μg/ml BSA Fraction V (Gibco, 15260), and the following small molecules and cytokines: 1 μM PD0325901 (Stemgent, 04-0006), 1 μM IM-12 (Enzo, BML-WN102), 0.5 μM SB590885 (Tocris, 2650), 1 μM WH4-023 (A Chemtek, H620061), 10 μM Y-27632 (Stemgent, 04-0012), 20 ng/mL recombinant human LIF (PeproTech, 300-05), and 10 ng/mL Activin A (Peprotech, 120-14)] were dissociated with TrypLE for 5 minutes at 37°C. Naïve hPSCs were dissociated to single cells with TrypLE and manual disruption, the enzyme was neutralized with fibroblast medium [DMEM (Millipore Sigma, #SLM-021-B) supplemented with 10% FBS (Gibco 26140079), 1X GlutaMAX (Gibco, 35050), and 1% penicillin-streptomycin (Gibco, 15140)], and cells were centrifuged at 1,200 rpm for 3 minutes. The supernatant was removed, and cells were washed once with DMEM/F12 and centrifuged again at 1,200 rpm for 3 minutes. The cells were resuspended in 1.5 mL of 5i/L/A medium and mitomycin C-inactivated mouse embryonic fibroblasts were removed from the cell suspension by incubation on gelatin-coated plates for 45 minutes at 37°C 5% CO_2_. The cells remaining in suspension were collected and counted.

Aggrewells were coated with anti-adherence solution (StemCell Technologies) for 10 minutes at room temperature. The wells were washed once in DMEM/F12, then 1 mL of 5i/L/A medium was slowly added to each well. A single cell suspension of 100,000 cells per 500 μL was added to each well, such that the total medium was 1.5 mL per well. The plate was centrifuged at 1,000 rpm for 2 minutes to ensure an equal distribution of cells per microwell. The cells were placed back in a 37°C, 5% CO_2_, and 5% O_2_ incubator for one day. On the following day, the medium was changed to N2B27 with Sodium Pyruvate (NaPy) without antibiotics [1:1 DMEM/F12:Neurobasal (Gibco), 1x N2 100X supplement (Gibco, 17502), 1x B27 50X supplement (Gibco, 17504), 1X GlutaMAX (Gibco 35050), 1X MEM NEAA (Gibco, 11140), 0.1 mM β-mercaptoethanol (Millipore Sigma, 8.05740), 1mM NaPy (Corning 25-000 CI)]. 1 mL of N2B27+NaPy medium was changed twice per well to wash out the 5i/L/A medium. On the third day, N2B27+NaPy medium was changed to blastoid induction medium (BIM) [1:1 DMEM/F12: Neurobasal, 0.5x N2 100X supplement (Gibco, 17502), 0.5x B27 50X supplement (Gibco, 17504), 0.5x GlutaMAX, 0.5x MEM NEAA (Gibco, 11140), 1mM NaPy (Corning 25-000 CI), 0.1 mM β-mercaptoethanol (Millipore Sigma, M3148), 0.5% ITS-X (Gibco 51500-056), 0.5% Knock-out Serum Replacement (Gibco 10828028), 0.1% FBS (Cytiva, SH30088.03), 1μM PD0325901 (Tocris 41-9210), 1 μM A83-01 (Tocris TB2939RMU10), 0.5 μM WH-4-023 (A Chemtek S1180), 0.25 μM IM-12 (Enzo BML-WN102), 25 ng/mL rhEGF (Qkine Qk011-0100), 3 μg/mL Ascorbic Acid (FujiFilm 013-12061), and 400 μg/mL Valproic acid (Sigma Aldrich P4543)]. 1 mL of BIM was changed twice per well to wash out the N2B27+NaPy medium. On each following day, 1 mL of fresh BIM was exchanged without perturbing the blastoids. Collection of samples was performed on D9 when most blastoid cavitation was observed.

### Single-cell Dissociation, Collection, Small-Seq and Co-Seq Library Preparation

Mouse embryos spanning the 2-through 64-cell stages and day 9 blastoids were dissociated into single cells and collected as we have previously described^117^. Individual cells, oocytes, or 0.1 µL aliquots of purified sperm samples (approximately 6300-8700 individual sperm cells per mouse) were then dispensed directly into 3 µL of Small-Seq lysis buffer^23^ and stored in a -80°C freezer. Following thawing and cell lysis, Small-Seq library preparation was performed as previously described^14,23^. Mouse libraries were sequenced on the NextSeq 550 (High output kit, Illumina) and blastoid libraries were sequenced on the NextSeq 2000 (High output kit, Illumina) To determine which cell type sncRNA profiles correspond to, dissociated mouse embryos (2-cell through 64-cell stages) and and day 9 blastoids were dispensed directly into 4 uL of Co-Seq Lysis buffer for both Small-Seq^23^ and Smart-Seq2^122^ library preparation, as we have previously described in detail^14^. Both the sncRNA (Small-Seq) and the mRNA (Smart-Seq2) portions were sequenced with the NextSeq550 (High output kit, Illumina)^25^.

### Pre-processing and quality control of the mRNA portion of Co-seq data

For the mRNA expression profiles of single-cell mouse Co-Seq data, sequenced reads were aligned to the mouse reference genome mm10 (v2020-A, obtained from the 10x Genomics website) using STAR (v2.5.3a) with default parameters^123^. Raw read counts were quantified by rsem-calculate-expression from RSEM (v1.3.0)^124^, using the option “--single-cell-prior”. Low-quality cells were removed by applying a threshold of at least 1000 detected genes (nGene > 1000) and less than 25% mitochondrial gene expression (percent.mito < 25%), resulting in 313 retained cells. Gene expression counts were normalized to library size after excluding mitochondrial genes. For the mRNA expression profiles of single-cell blastoid Co-Seq data, reads were processed using the same tools as mouse data but aligned to the human reference genome GRCh38 (v3.0.0, obtained from the 10x Genomics website). Low-quality cells were filtered using thresholds of nGene > 1500, percent.mito < 25% and uniquely mapping ratio more than 25%, leaving 100 high-quality cells for downstream analysis. The sex of each cell and mouse embryo was inferred based on the expression of Y-linked genes, as previously described^2^. Cells with a total Y chromosome expression (∑RPKM ChrY) greater than 75 were classified as male, while those with ∑RPKM ChrY less than 25 were classified as female. Embryo sex was determined according to the ∑RPKM ChrY values expressed in the majority of cells from the same embryo. No sex-specific differences were observed across lineages during mouse preimplantation development. The only potential difference between female and male ICM cells was miR-421-3p, which showed higher expression in female ICM cells (log2FC = 3.89, FDR = 0.0565), but no difference between female and male TE cells. Therefore, all data was collapsed to not consider sex for all downstream analyses.

### Integration of mRNA portion of Co-Seq data with embryonic reference data

To determine the cell types within our Co-Seq dataset, we processed the mouse and human mRNA libraries separately. For the mouse mRNA portion, we integrated our mRNA data with a previously published scRNA-seq dataset^36^ using canonical correlation analysis (CCA) as implemented in the Seurat R package (v5.1.0^125^). Integration was performed with 25 dimensions, 2,000 anchor features, and a k.filter setting to 100, using the functions “FindIntegrationAnchors” and “IntegrateData”. After integration, we applied principal component analysis (PCA) to the scaled data, followed by dimensionality reduction with Uniform Manifold Approximation and Projection (UMAP) using the “RunUMAP” function. Clustering was carried out with the “FindNeighbors” and “FindClusters” functions. Cell-type identities were assigned to clusters based on well-established lineage-specific gene signatures^36^. During integration, 88 cells with ambiguous annotations from Posfai et al.^36^ were excluded^126^.

For the human mRNA data, cell lineages were annotated by examining canonical lineage markers within clusters generated by the standard Seurat pipeline, using the top 2,000 variable genes and the top 20 principal components. Cell annotations were further validated with our Early Human Embryogenesis Prediction Tool^105^ (https://petropoulos-lanner-labs.clintec.ki.se/; Fig. 7c). Combined with the prediction results, the absence or very low expression of post-implantation lineage markers confirmed that potential post-implantation-like lineages were not present (Extended Data Fig. 9a).

### Pre-processing and quality control of Co-seq sncRNA data

Mouse sncRNA reads were processed using a modified version of the Small-Seq pipeline^14,23,25^. First, unique molecular identifiers (UMIs, 8 bp) were extracted from sequence reads and appended to read names using UMI-tools (v0.4.4)^127^. Adapters and poly(A) sequences were then trimmed with cutadapt (v1.17)^128^ using the parameters “-e 0.1 -o 3 -m 18 -M 69 -u 2”. The “-M 69” parameter was chosen because the maximum read length in our small RNA sequencing libraries was 84 bp. To ensure that retained reads originated from short transcripts, at least 5 bp of adapter or poly(A) sequence trimming was required.

Clean reads were aligned to the mouse mm10-2020-A reference genome using Bowtie (v1.0.0) with parameters “-a --best --strata -v 2 -m 50 -S -q -p 2“^129^. To minimize false-positive alignments, reads shorter than 20 bp with ≥1 mismatch and reads 20–40 bp with ≥2 mismatches were excluded. Mismatches at the terminal base were disregarded, accounting for the “CCA” addition during tRNA maturation and miRNA modification dynamics. Unmapped reads were subjected to recursive soft clipping from the 3′ end, trimming one nucleotide at a time and remapping with Bowtie (“-a --best --strata -v 1 -m 50”), until a minimum length of 3 nt was reached; the same procedure was then applied to the 5′ end. Reads with final mapping lengths shorter than 17 bp were discarded. PCR duplicates were collapsed, and RNA molecules were counted with UMI-tools using the “dedup --method directional” option. Deduplicated soft-clipped reads were remapped to the genome using the same procedure to recover multi-mapping information potentially lost during deduplication. Reads were annotated against multiple reference databases: miRBase v22.1^130^ (miRNA), Gencode V38^131^ (snoRNA, snRNA, rRNA, and others), GtRNAdb V19^132^ (tRNA), and piRBase V3.0^133^ (piRNA). The reads were assigned to each category in a hierarchical manner based on their overlap with the above annotations in the same strand with priority assigned in the following order: miRNA, rRNA, snoRNA, snRNA, tRNA, piRNA, and other transcripts. Reads overlapping mature miRNAs and piRNAs were additionally required to be less than 40 nt.

Cell-level quality control was performed based on three criteria: (1) a minimum of 0.35 million sequenced reads per cell; (2) less than 10% of UMIs derived from mitochondrial RNAs; and (3) >100 expressed miRNA molecules per cell. Co-Seq–derived sncRNA data were used exclusively for data integration and correlation analysis with gene expression, and excluded from downstream analyses to minimize batch effects from library preparation and reduce potential technical artifacts.

Human sncRNA reads were processed using the same pipeline, as we previously described in our published human embryonic paper^14^. For cutadapt, the “-M” parameter was set to 69 for Small-Seq human blastoids data and 81 for the sncRNA portion of Co-seq human blastoids data, reflecting differences in sequencing read length.

### miRNA expression analysis of Co-Seq data

For the sncRNA portion of Co-Seq data from mouse and human blastoids, miRNAs were retained if they showed at least one count in a minimum of two cells. Log-normalized counts were calculated using the deconvolution strategy implemented in the computeSumFactors function of the scran package (v1.14.6)^134^. Dimensionality reduction was performed with the Seurat package, using the top 150 most variable miRNAs. The top 10 principal components were then computed with function “RunPCA” and used for UMAP-based dimensionality reduction and clustering via function “RunUMAP” and “FindClusters”, respectively. To assign cluster identities based on miRNA expression, we leveraged indicator cells from the mRNA portion of the Co-Seq data, designating them as either mouse blastocyst lineages or blastoid epiblast-like (ELCs) and trophectoderm-like cells (TLCs). Cells with conflicting annotations between the mRNA-based and miRNA-based clustering were labeled as “unknown” and excluded in the integration with full-cell Small-Seq data.

### Pre-processing, quality control and normalization of Small-Seq data

Read mapping, feature counting and normalisation for mouse embryonic and human blastoids full-cell Small-seq data were using the same procedures as those employed for the Small-seq portion of the Co-Seq data.

### Integrated analysis of Small-Seq datasets and Co-Seq datasets

For mouse embryonic data, subtle batch differences were observed between Co-Seq and Small-Seq. Therefore, we combined Co-Seq and Small-Seq cells and performed normalization using the “computeSumFactors” function in the scran package. Standard Seurat processing was then applied, using the top 250 most variable miRNAs. Subsequently, the top 25 principal components were computed with “RunPCA” and used for UMAP-based dimensionality reduction and clustering via “RunUMAP” and “FindClusters”, respectively. Dimensional reduction was conducted separately for miRNA, tRNA, piRNA, snRNA, rRNA and snoRNA expression. For the human blastoid data, noticeable batch differences were observed between Co-Seq and Small-Seq, potentially due to differences in sequencing read length and platform. To integrate the two datasets, we first applied rescaled normalization using the multiBatchNorm function from the batchelor package^135^, ensuring expression levels were comparable across datasets. We then performed integration with the CCA-based approach implemented in Seurat, using 20 dimensions, 250 anchor features, and a k.filter value of 50. The final assignment of cell identity for the formed clusters was determined by leveraging the miRNA signatures obtained from the Co-Seq dataset with known lineages. For dimensionality reduction based on tRNA, piRNA, snRNA, rRNA, and snoRNA expression in full-length Small-Seq data from human blastoids, we applied the standard Seurat workflow, following the same procedures and parameter setting as used for the sncRNA portion of the Co-Seq data. Integration between human blastoids and human embryonic data^14^ was performed using the CCA-based approach implemented in Seurat, with 20 dimensions, 250 anchor features, and both k.filter and k.weight set to 50. “unclassified clusters” from human embryonic cells were excluded. Small-Seq data of Naive and Primed cells were downloaded from Faridani et al^23^ and reprocessed as we previously described^14^.

### Marker miRNA detection and differential miRNA expression analysis

Final miRNA markers for each cluster were identified using the “FindAllMarkers” function in the Seurat package. To define blastocyst (ICM & TE) and prelineage markers, ICM and TE cells were grouped together, and 2C to morula cells were combined. Markers with adjusted p-values (p_val_adj < 0.05) were retained (Supplementary Data 2). Visualization of marker miRNA loci was performed using the circlize R package (v0.4.16)^136^. Normalization size factors for merged reads used in IGV^137^ were calculated with the computeSumFactors function in the scran package.

Differential miRNA expression analysis across lineages and developmental stages was performed using the two-sided Wilcoxon test included in “FindMarkers” function. False discovery rates (FDRs) were calculated with the Benjamini–Hochberg method. miRNAs with FDR < 0.05 and log2 fold change > 0.1 were considered significant. Considering the batch effects among human embryos, blastoids, and stem cells, differentially expressed miRNAs between ICM vs ELC, ICM vs TLC, and TE vs TLC were further filtered by requiring expression of an individual miRNA to be in at least 50% of cells within the up-regulated groups and absence of expression in at least 90% of cells within the down-regulated groups, based on normalized values obtained from the multiBatchNorm function.

### Potential inheritance of sncRNAs

We first quantified all UMIs for sncRNAs expressed in the individual cell types (Extended Data Fig. 8a). To determine exclusive expression among the cell types, we applied a cutoff that “25% of cells with expression” to reduce the likelihood that undetected sncRNAs were simply due to low expression frequency. Among the sncRNAs detected with UMIs in sperm, oocyte, or 2-cell embryos, we classified them into the following categories. sncRNAs with detectable UMI reads in sperm, oocyte, and 2-cell embryos were classified as expressed in all three cell types and termed ‘shared’. sncRNAs expressed in at least 25% of sperm cells but with no UMI reads in either oocyte or 2-cell embryos were classified as “only expressed in sperm”. The same definition applies to our other categories such as “only expressed in oocyte” and “only expressed in 2-cell embryos”. sncRNAs expressed in at least 25% of both sperm and oocyte cells, but absent (no UMIs detected for individual sncRNA) in 2-cell embryos, were classified as “expressed in sperm and oocyte but not in 2-cell embryo”. The same definition applies to “expressed in sperm and 2-cell embryo but not in oocyte” and “expressed in oocyte and 2-cell embryo but not in sperm”. To avoid false positives, sncRNAs that do not fall into any of the above categories were assigned to the “uncertain” group.

### Pseudotime reconstruction and trajectory inference

Single-cell pseudotime trajectories were computed using the R package slingshot (v2.10.0), which infers lineage structures in a low-dimensional space^138^. Briefly, pre-computed cell embeddings and lineage annotations derived from miRNA expression profiles of embryonic cells were used to construct the UMAP reduction generated by the standard Seurat workflow based on the top 100 most variable miRNAs and the top 10 principal components. Subsequently, pseudotime trajectories were constructed by utilizing the function “slingshot”, with setting the start cluster to “2C&4C”. Individual pseudotimes were then inferred using the function “slingPseudotime” with the parameter “na = TRUE”. Since pseudotimes were computed separately for the ICM and TE trajectories, we aligned the two trajectories through the following rescaling procedure:

~~~
scale.factor = mean(Psd_ICM_max_64C - Psd_prelineages_max) / mean(Psd_TE_max_64C
- Psd_prelineages_max)
Psd_TE_mod = (Psd_TE - Psd_prelineages_max) * scale.factor + Psd_prelineages_max
~~~

Here, Psd_TE_max_64C represents the latest pseudotime for 64-cell TE cells, Psd_ICM_max_64C represents the latest pseudotime for 64-cell ICM cells, and Psd_prelineages_max represents the latest pseudotime for prelineage cells. Psd_TE corresponds to the individual pseudotime values for TE cells greater than Psd_prelineages_max, and Psd_TE_mod denotes the rescaled pseudotime values for TE cells.

To visualize individual pseudotimes, principal curves were fitted to the Seurat object. Trajectory-associated miRNAs were selected using function DynamicHeatmap from the R package SCP (v0.5.6, https://github.com/zhanghao-njmu/SCP), with the added requirement of having differential expression detected either between adjacent stages or between each later stage and the 2C stage. These analyses were performed separately for the ICM and TE lineages. Finally, expression patterns of trajectory-related miRNAs were further examined using k-means clustering implemented in the R package pheatmap (v1.0.12)^139^.

### Weighted (miRNA) gene co-expression network analysis (WGCNA)

To identify and analyze miRNA with similar expression patterns in theSmall-Seq data from whole cells, we applied hdWGCNA (v0.4.04)^63^. Transcriptionally similar cells from the same lineages were grouped using the “MetacellsByGroups” function (k=15, min_cells=10). The “TestSoftPowers” function was then used to determine the optimal soft-thresholding power that produces a scale-free gene co-expression network, followed by network construction with the “ConstructNetwork” function (minModuleSize=5). To leverage Co-Seq data for assessing miRNA–gene expression correlations, the previously built co-expression network from the Small-Seq data was projected onto the miRNA portion of the Co-Seq dataset using the “ProjectModules” function. Module robustness and preservation were assessed with the “ModulePreservation” function, evaluated by 200 permutations within both the Small-Seq and Co-Seq datasets, and summarized using standardized Z-scores of quality and preservation. Additionally, a gene co-expression network was also constructed from the mRNA portion of the Co-Seq data using the same function as for miRNA with default setting for “ConstructNetwork” function.The expression profile of each module was summarized by its module eigengene (ME), calculated with the GetME function. Module hubness, defined as the module eigengene-based connectivity (kME), was ranked by the correlation between miRNA (or gene) expression and the corresponding ME using the “GetModules” function. Differential module eigengene analysis was performed with the “FindDMEs” function using a two-sided Wilcoxon test, and modules with p < 0.05 were considered significant. The ModuleRadarPlot and ModuleFeaturePlotfunction were used to visualize module expression, and hub miRNAs were visualized using the “HubGeneNetworkPlot” function. All above functions mentioned are from hdWGCNA R package.

### Analysis of mRNA and target gene correlations

Candidate miRNA–mRNA target relationships were obtained from miRDB (v6.0)^64^ and TargetScan (vert_80)^65^, and combined into a union set. Spearman correlation coefficients between miRNA expression and their predicted target gene expression were calculated using all Co-Seq cells with consistent miRNA and gene annotations, or using blastocyst cells when the miRNA modules were identified as differentially expressed between ICM and TE. One-sided Student’s t-tests were applied to assess positive or negative correlations. Given that miRNA and mRNA expression are typically inversely related^140,141^, only significant negative correlations (p < 0.05) were displayed in the figures, while all significant correlations were reported in the supplementary data (Supplementary Data 6). Pairwise Spearman correlations between miRNA modules and gene modules were also computed. Significant module–module correlations were required to meet two criteria: (1) p < 0.05 with correlation < –0.2, and (2) support from at least one significant miRNA–gene pair within the corresponding modules.

### miRNA target and transcriptional factor regulatory enrichment analysis

To assess whether miRNA target genes were enriched in specific gene modules, one-sided Fisher’s exact tests were performed, using all miRNAs with annotated target genes as the background. Similarly, to evaluate whether transcription factor (TF) regulatory interactions were enriched in specific miRNA modules, one-sided Fisher’s exact tests were also applied. Predicted TF–miRNA regulatory interactions were obtained from TransmiR(v3.0), and TFs with recorded miRNA regulation were used as the background. Significant enrichment was defined as odds ratio (OR) > 1 and p-value < 0.05.

### Pathway enrichment analysis of gene modules

Pathway enrichment analysis of each gene module was performed using the enricher function from the clusterProfiler package^142^. WikiPathway annotations and gene sets were obtained from the Molecular Signatures Database^143^ and the Enrichr database^144^. Significantly enriched WikiPathways were defined as those with p-value < 0.05.

### Cross-species miRNA expression comparison between human and mouse

Conserved miRNAs between human and mouse were defined with two inclusion criteria: miRNAs with identical mature sequences in both species and miRNAs with the same name and identical seed sequence (positions 2–8). miRNA families were regarded as conserved if they shared the same family name in both species and contained at least one member with an identical seed sequence. Individual miRNAs belonging to these shared families but lacking sequence conservation were defined as species-specific members of conserved families. All other non-conserved miRNAs were classified only as species-specific miRNAs. All DE miRNAs (union set) across lineages and developmental stages in human and mouse preimplantation embryos were selected. Among these, conserved miRNAs with similar expression patterns were identified by exhibiting the highest expression in the corresponding lineages or developmental stages in both species.

### Immunofluorescence Analysis and Confocal Microscopy

For analysis of protein expression via immunofluorescence and one photon confocal microscopy, embryos were fixed, permeabilized, blocked, and incubated overnight at 4°C with the following primary antibodies (Methods Table 1) as we have previously described^14^. Subsequently, embryos were incubated with secondary antibodies (Methods Table 1) for 2h at RT, before finally being placed in a 1:10000 dilution of Hoechst nuclear stain (Invitrogen, cat # H1399) for 10 min at RT. All images were acquired using an Olympus FV1000 upright microscope (Olympus, Japan), equipped with a XLUM Plan FL N 20x/ 1.00 Water objective, as previously described^14^.

Images were analysed using ImageJ software version 2.14.0/1.54 f to assess blastocyst formation, lineage cell counts, and to quantify the relative protein expression of genes of interest. Nuclear fluorescence intensity (arbitrary fluorescence units) measurements from each cell in all channels were normalised to the average expression of Hoechst (nuclear stain) within the corresponding embryo. Statistical analysis was performed using GraphPad Prism 4 statistical software. All values were reported as means ± standard error. Differences between control and treated groups were analysed using the two-sided Student’s t test or Mann Whitney test where appropriate. P < 0.05 was considered a statistically significant difference.

## Methods Tables

**Table 1.** Primary and secondary antibodies used in immunofluorescence experiments.

| Antibody | Source | Catalog # | Species | Dilution |
| --- | --- | --- | --- | --- |
| Anti-Active Yap | Abcam | ab205270 | Rabbit | 1:100 |
| Anti-Sox2 | eBioscience | 14981182 | Rat | 1:100 |
| Anti-Cdx2 | Biogenex | MU392A-5UC | Mouse | 1:250 |
| Anti-Rabbit 488 | Life Technologies | A21206 | Donkey | 1:1000 |
| Anti-Rat 555 | Life Technologies | A48270 | Donkey | 1:1000 |
| Anti-Mouse 647 | Life Technologies | A31571 | Donkey | 1:1000 |

## Author Contributions

S.P. conceived the study and supervised the project. S.B. and S.P. designed and S.B. performed the experiments. C.Z. performed the bioinformatic analysis. S.B., S.P. and C.Z. analysed the data, interpreted the results and wrote the manuscript. R.Y. and T.W.T. generated the human blastoids, which S.B. then collected and prepared libraries from and data were analyzed by C.Z.

## Acknowledgements

This work was supported by funding from The Canadian Institutes of Health Research (PJT-178082, SP), the Swedish Research Council (2016-01919, SP), Canadian Foundations for Innovation-John R. Evans Leader (38876, SP), Fonds de recherche du Québec - Santé (SB) and by the Université de Montréal Bourse de la Réussite étudiante aux cycles supérieurs (SB). SP holds the Canadian Institutes of Health Research Tier 2 Canada Research Chair in Functional Genomics of Reproduction and Development (950-233204). The computations and data handling were enabled by resources in project NAISS 2023/22-988 and NAISS 2023/23-490 provided by the National Academic Infrastructure for Supercomputing in Sweden (NAISS), partially funded by the Swedish Research Council through grant agreement no. 2022-06725. We would also like to thank Dr. Christine LaFleur and the Sarah Kimmins lab with assistance obtaining mouse sperm, and the CRCHUM Cellular Imaging Core Facility for their equipment and expertise with imaging procedures. Blastoid experiments in the Theunissen lab were funded by a Mallinckrodt Scholar Award from the Edward Mallinckrodt, Jr. Foundation to T.W.T.

## Extended Data Figures

**Extended Data Fig. 1.**
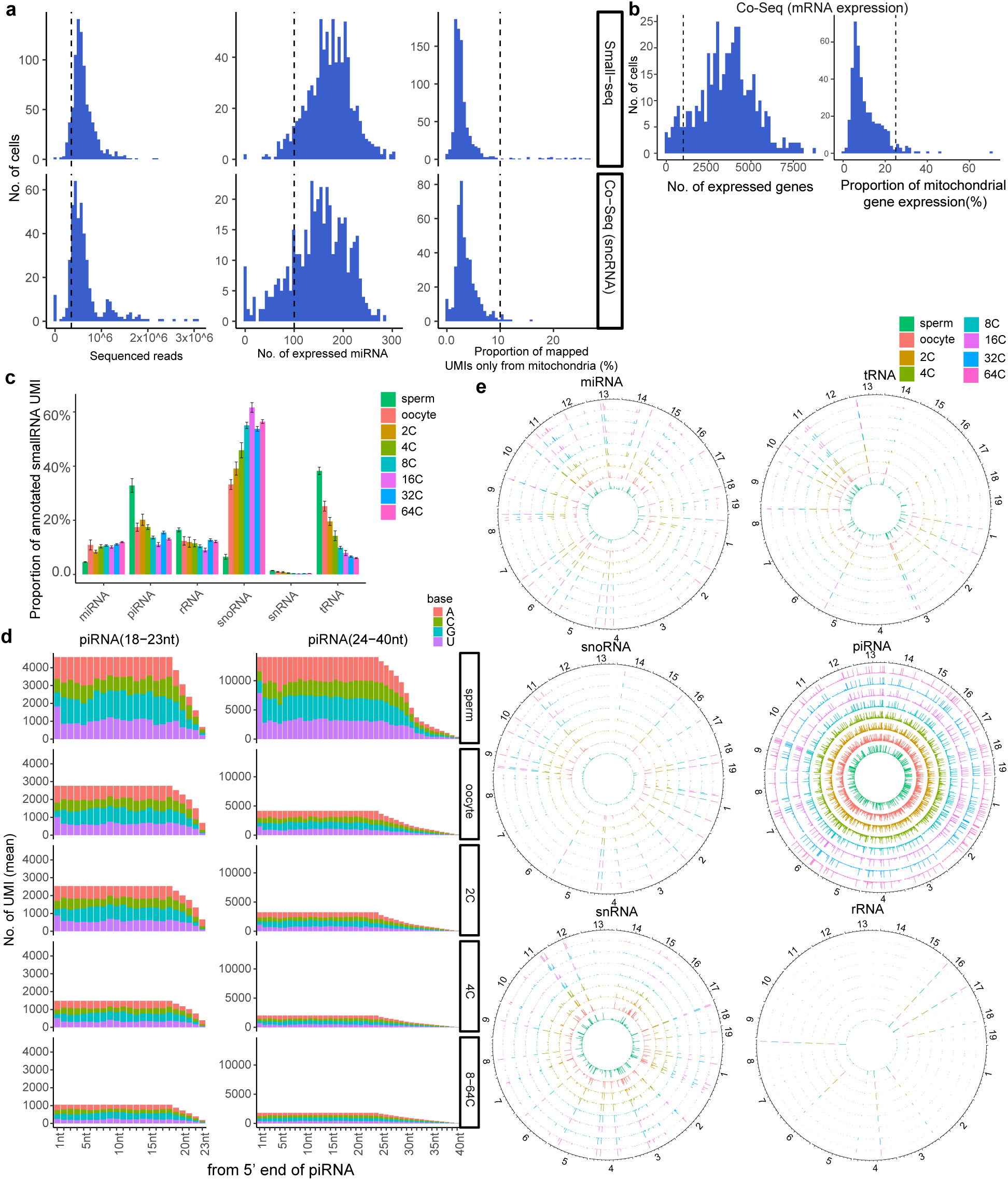
Quality control of mouse preimplantation Small-Seq and Co-Seq data. **a,** Distribution of sequenced reads, number of expressed miRNAs, and proportion of UMIs uniquely mapped to mitochondria in the sncRNA part of Co-Seq and Small-Seq. The dashed line indicates the cutoff used to remove low-quality samples. **b,** Distribution of number of expressed genes, and proportion of mitochondria gene expression in the mRNA part of Co-Seq. The dashed line indicates the cutoff used to remove low-quality samples.**c,** proportion of UMIs annotated to microRNA (miRNA), small nucleolar RNA (snoRNA), small nuclear RNA (snRNA), transfer RNA (tRNA), ribosomal RNA (rRNA), and piwi-interacting RNA (piRNA) by day of embryonic development. The boxplot rectangles represent the first and third quartiles, a horizontal line inside the box indicates the median value. **d**, Nucleotide distribution starting from the 5′ end of piRNAs, with ‘18–23 nt’ and ‘24–40 nt’ representing UMI read lengths range. **e,** Circular plots showing the genomic distribution of the sncRNA subtypes across the 19 autosomal chromosomes.

**Extended Data Fig. 2.**
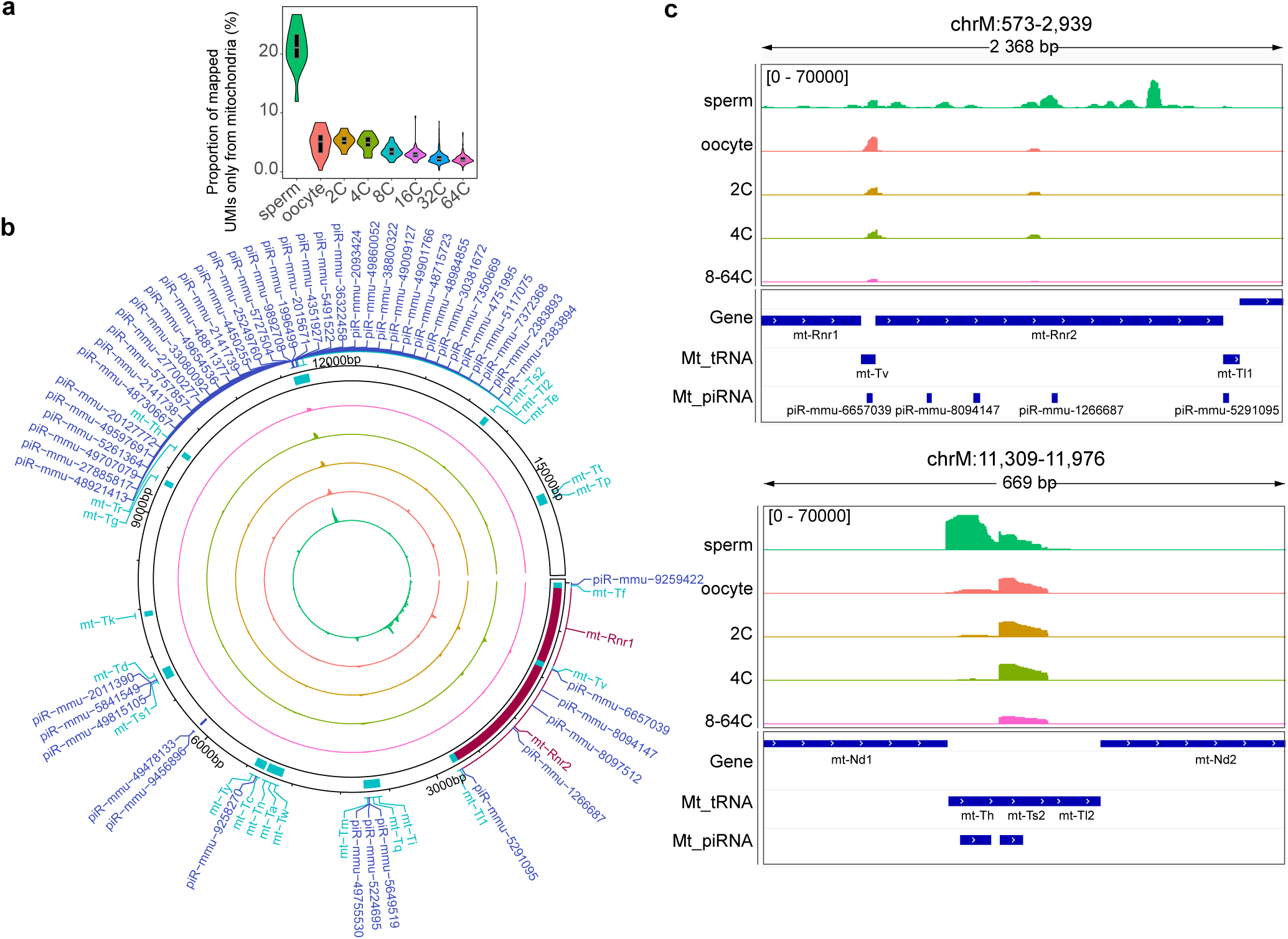
Mapping of sncRNAs to the mouse mitochondrial genome during preimplantation development. **a,** Percentage of total reads mapped to mitochondrial (Mt) DNA. **b,** Circular plot showing the genomic distribution of reads mapped to mtDNA. **c,** Regions where the majority of reads mapped to the mtDNA in sperm are localized.

**Extended Data Fig. 3.**
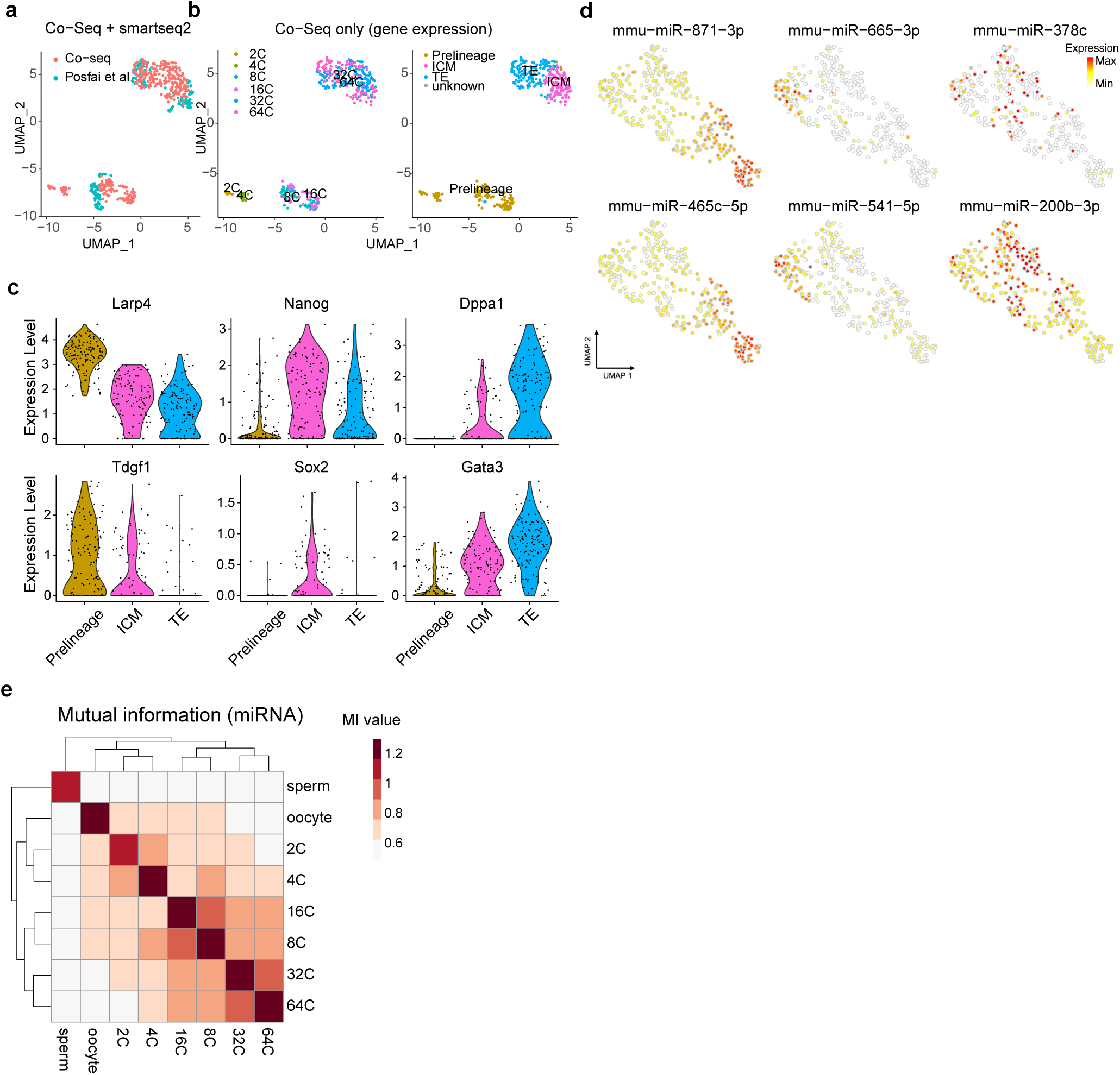
Lineage identification and miRNA dynamics across mouse embryo development continued. **a,** Uniform Manifold Approximation and Projection (UMAP) showing the integration of Co-Seq data (mRNA portion) with Posfai et al to establish cell identities based on gene expression. **b**, UMAP including only the Co-Seq data from the integration in (a), colored by developmental timepoint and lineage annotation. **c**, Violin plots shown lineage marker gene expression in Co-Seq cells. **d**, Expression of selected lineage-specific miRNA markers in the sncRNA portion of Co-Seq data, prelineage: miR-871-3p, miR-465c-5p; ICM: miR-665-3p, miR-541-3p; TE: miR-378c, miR-200b-3p. **e,** Mutual information among different embryonic stages based on miRNA expression data using full-cell Small-Seq dataset.

**Extended Data Fig. 4.**
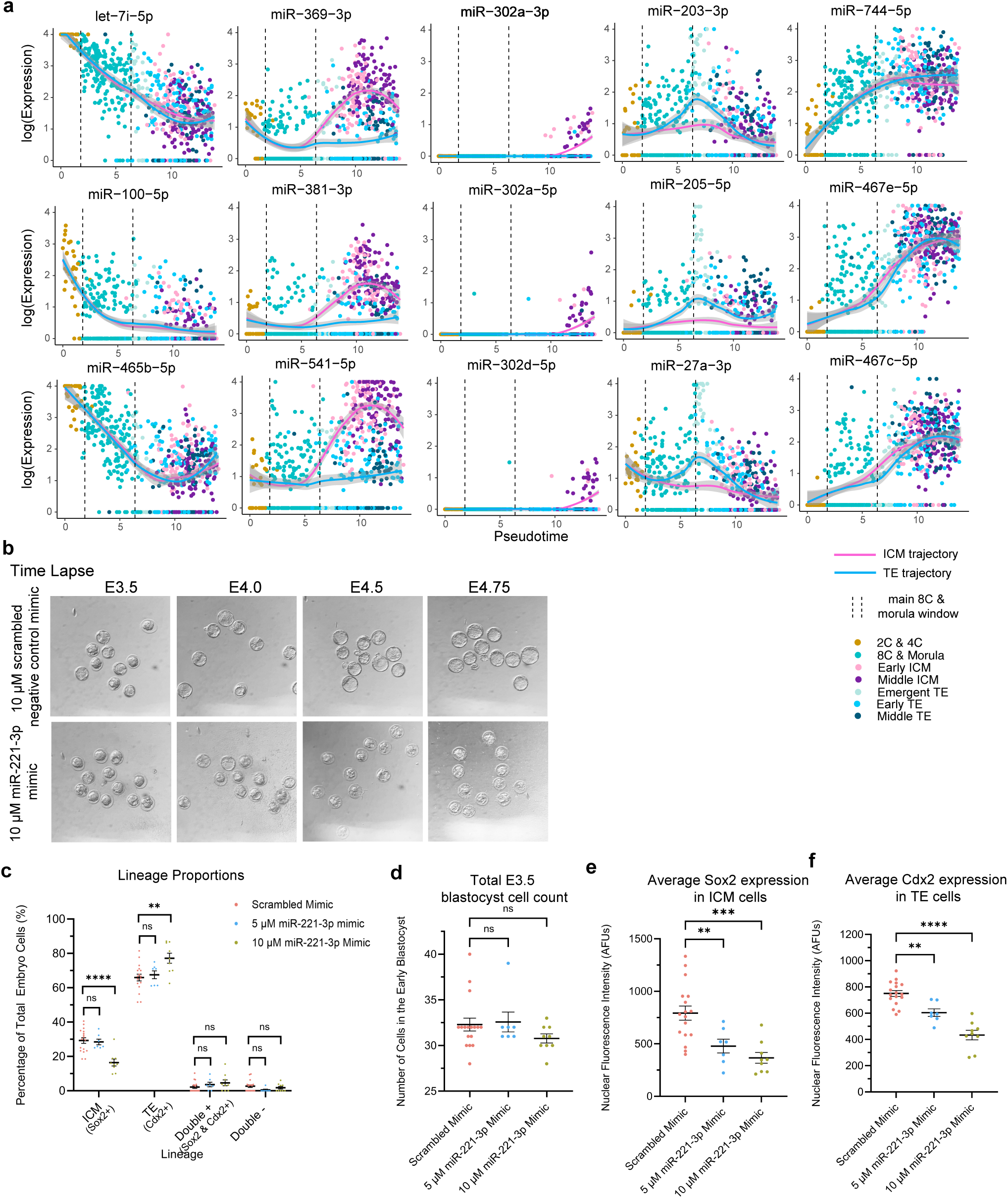
Continuation of lineage-specific pseudotime analysis and miR-221-3p mimic treatment. **a,** Expression of several key miRNAs across pseudotime associated with pre-lineage cells (first column), ICM trajectory (second and third columns), TE trajectory (fourth column) and blastocyst formation (fifth column). The confidence interval (error bands, 95%) is indicated by bandwidth. The measure of centre and confidence intervals were calculated using the “loess” function with default parameters in R software. **b,** Representative images of time lapse experiment of blastocyst development when embryos are treated with miR-221-3p mimic at the 2-cell stage, all images taken at 13x magnification. **c,** Analysis of cell lineage allocation in early (32-cell) blastocysts treated with scrambled control versus miR-221-3p mimic, where each data point represents the proportion of total embryonic cells belonging to the ICM or TE in a single embryo. Lineage was assigned based on the exclusive expression of known lineage markers Sox2 (ICM) and Cdx2 (TE) using immunofluorescence and confocal microscopy. Double positive (+) cells expressed both Sox2 and Cdx2. Double negative (-) cells expressed neither Sox2 or Cdx2. **d,** Average number of embryonic cells in control versus miR-221-3p mimic-treated blastocysts, where each datapoint represents the total number of cells in one embryo. Quantified average nuclear fluorescence intensity (protein expression, measured in arbitrary fluorescence units (AFUs)) of **e,** Sox2 (in ICM cells) and **f,** Cdx2 (in TE cells) in early blastocysts treated with scrambled control versus miR-221-3p mimic. Each data point represents the average intensity of the cells in one embryo. Treatment/staining/imaging was performed in 4 separate batches. All statistical comparisons were conducted using either a One-Way ANOVA or non-parametric (Kruskal-Wallis) test where appropriate. Values are reported as means ± standard error. *ns* = *p* > 0.05, * = p < 0.05, ** = p < 0.01, *** = *p* < 0.001, **** = *p* < 0.0001.

**Extended Data Fig. 5.**
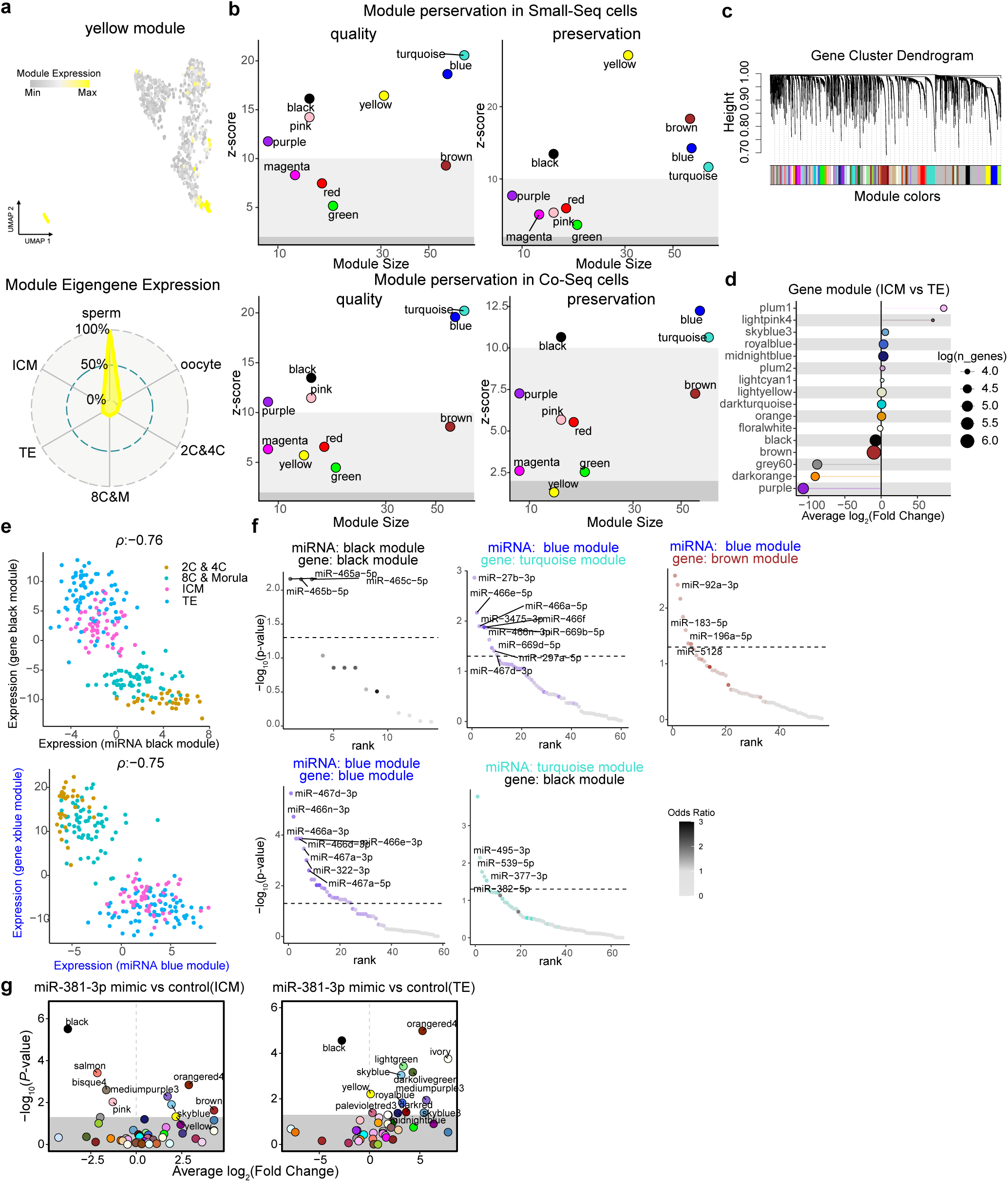
Additional interrogation of miRNA and gene module interactions. **a,** UMAP and radar plots of the yellow miRNA module, highly expressed specifically in mouse sperm samples. **b,** Preservation analysis of miRNA modules between the full-cell Small-Seq data and the sncRNA portion from the Co-Seq data. For each module, standardized Z-scores of quality from the preservation were calculated based on 200 permutations, using the full-cell Small-Seq data as the reference. Module preservation was considered strong if Z-summary > 10, weak to moderate if 2 < Z-summary < 10, and not supported if Z-summary ≤ 2. **c,** Cluster dendrogram detailing identification of gene modules in the gene portion of the Co-Seq data. Module colors are indicated at the bottom, with grey representing genes not assigned to any co-expression module. **d**, Significant differential module eigengenes (DMEs, P-value < 0.05) between the ICM and TE. The x-axis shows the average log₂ fold change (ICM vs. TE) for each gene module. Dot color corresponds to the assigned module, and dot size reflects the log-transformed number of genes within the module. **f**, Scatter plot showing the expression of eigengenes for black–black and blue–blue miRNA–gene modules using cells from the Co-Seq data. The Spearman correlation (ρ) is indicated at the top. **f**, miRNA target enrichment analysis showing the miRNAs with enriched targeting genes for each miRNA–gene module pair. The x-axis represents the top-ranked results sorted by one-sided p-values from Fisher’s exact test, and the color indicates the odds ratio from the same test. **g**, DMEs (P-value < 0.05) between miR-381-3p mimic and control cells in ICM and TE. Data reanalyzed from Russell & Zhao et al., 2024. Dot color indicates the assigned gene module.

**Extended Data Fig. 6.**
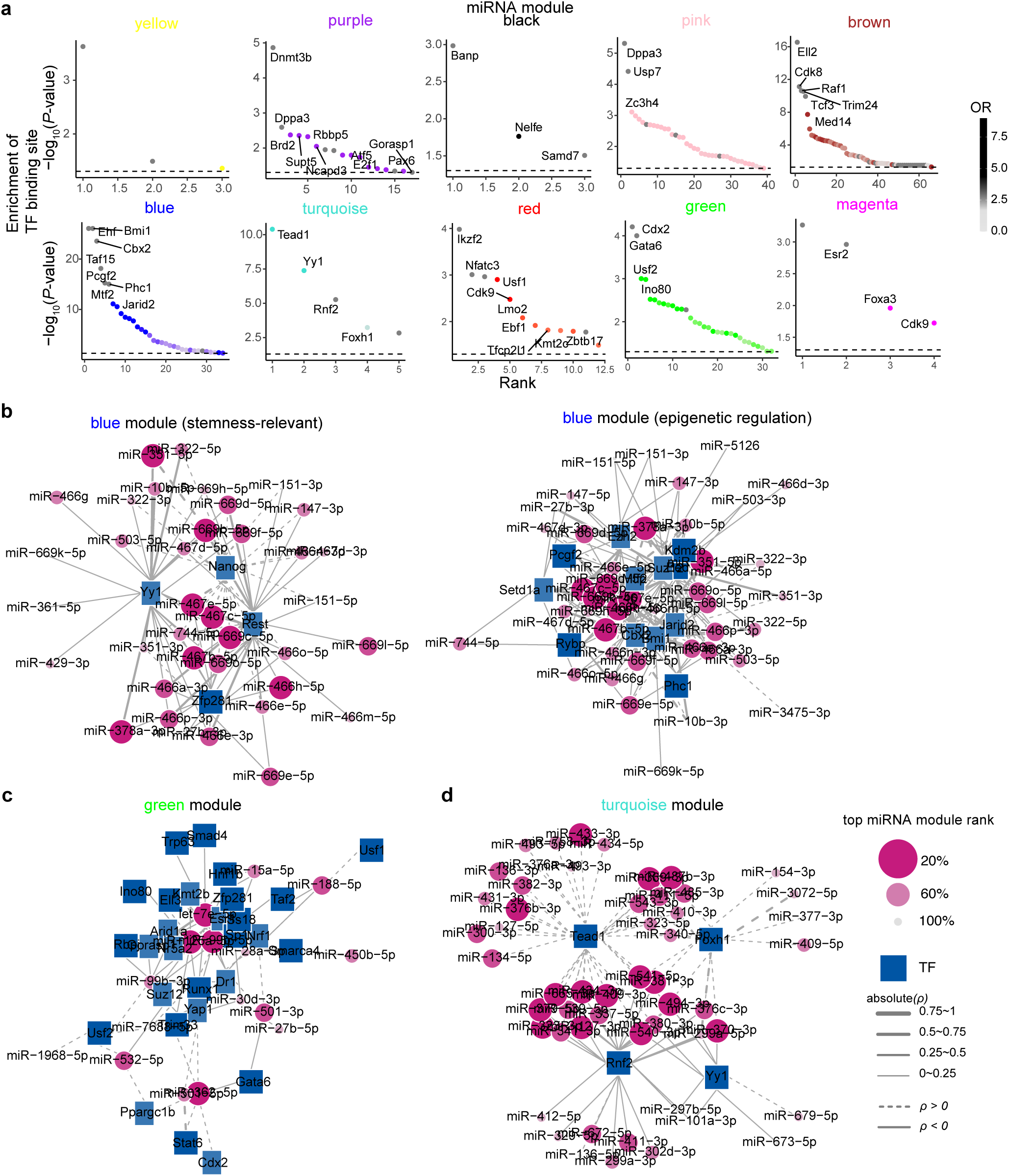
Transcription factor binding site enrichment analysis for mouse preimplantation embryo miRNA modules. **a,** Transcription factors (TF) with enriched promoter binding sites for each gene module. The x-axis represents the top-ranked results sorted by one-sided p-values from Fisher’s exact test, and the color denotes the odds ratio from the same test. **b-d,** Network plots showing TF with enriched promoter binding sites in the indicated miRNA modules. Only TF-miRNA pairs with significant Spearman correlations (p-value < 0.05) were included. Circles represent miRNAs and reflect their kME rank within the module. Line thickness indicates the Spearman correlation (ρ), and line type represents whether the correlation is positive or negative. Due to the complexity and function of TFs in the blue modules, the network plot for the blue miRNA modules was split into two parts (stemness and epigenetic regulation).

**Extended Data Fig. 7.**
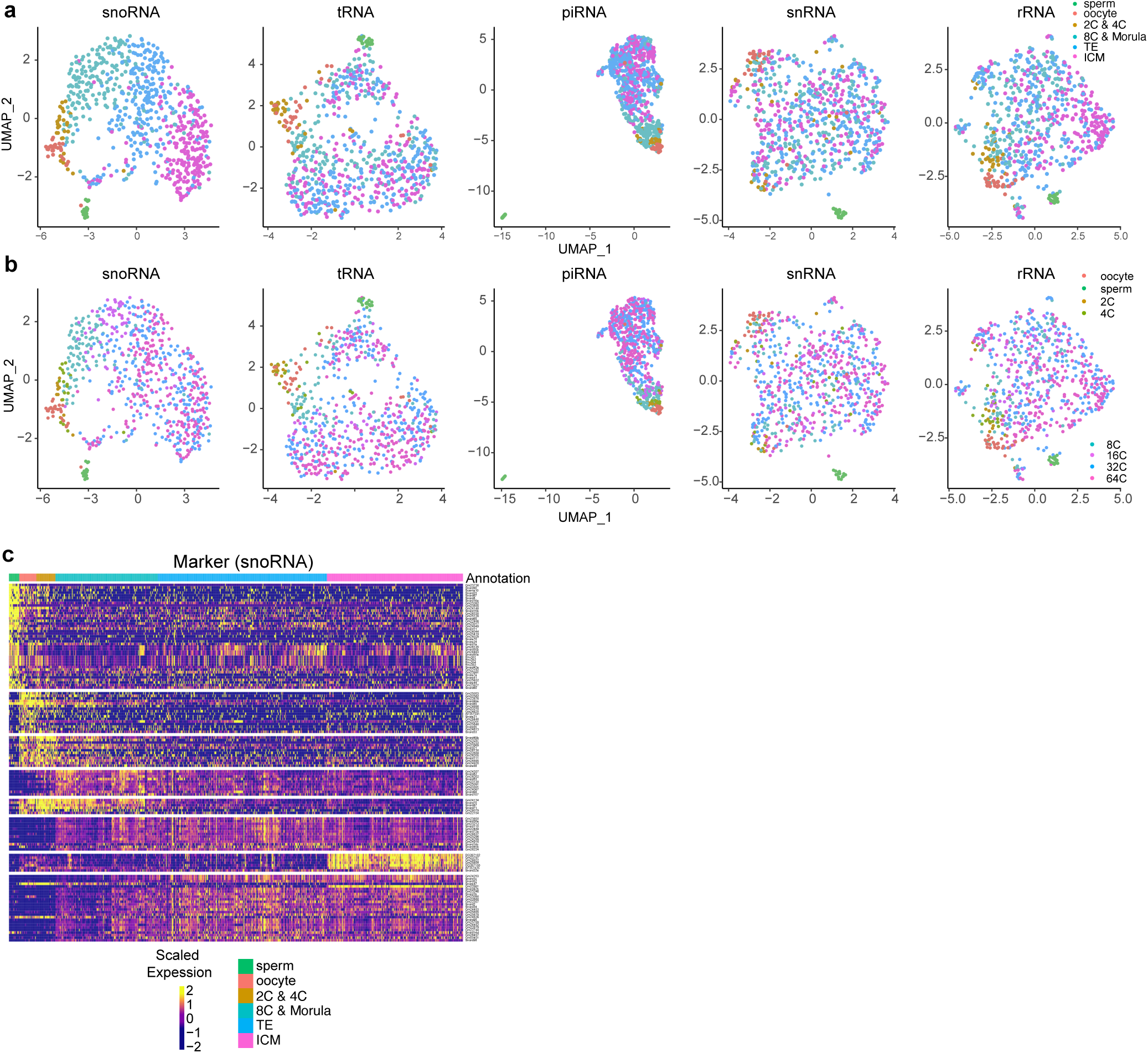
Expression dynamics of other sncRNA subtypes during mouse preimplantation development. UMAPs visualizing the single-cell expression profiles of the sncRNA subtypes labelled by **a,** lineage and **b,** developmental stage. **c,** Heatmap of top significantly enriched snoRNAs at each developmental stage and within each lineage.

**Extended Data Fig. 8.**
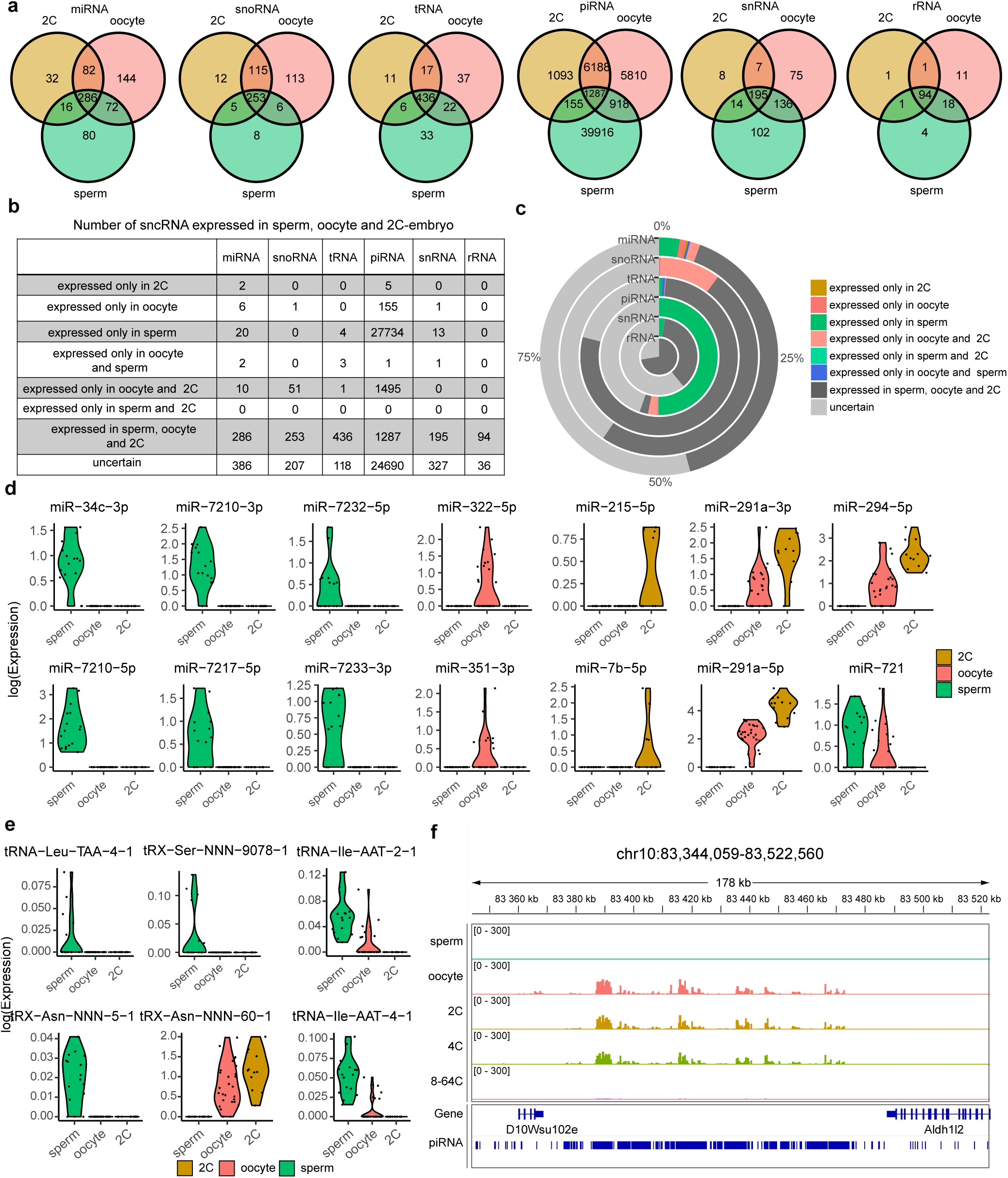
Analysis of sncRNA in sperm, oocytes, and 2-cell embryos. **a,** Venn diagram showing the overlap of sncRNA with UMI reads among sperm, oocyte and 2-cell-embryos. **b,** Table showing the number of sncRNA detected amongst sperm, oocyte, and 2-cell embryos. “Uncertain” indicates sncRNAs lacking UMI reads in specific lineages, possibly due to overall low expression frequency (% of cells expressing a specific UMI). **c**, Proportions of sncRNAs stratified by different categories. **d**-**e**, Violin plots showing log-normalized expression of selected miRNAs (**d**) and tRNAs (**e**). **f**, Integrative Genomics Viewer (IGV) showing the genomic coordinates and normalized reads coverage on chromosome 10. For a comprehensive list see Supplementary Data 7.

**Extended Data Fig. 9.**
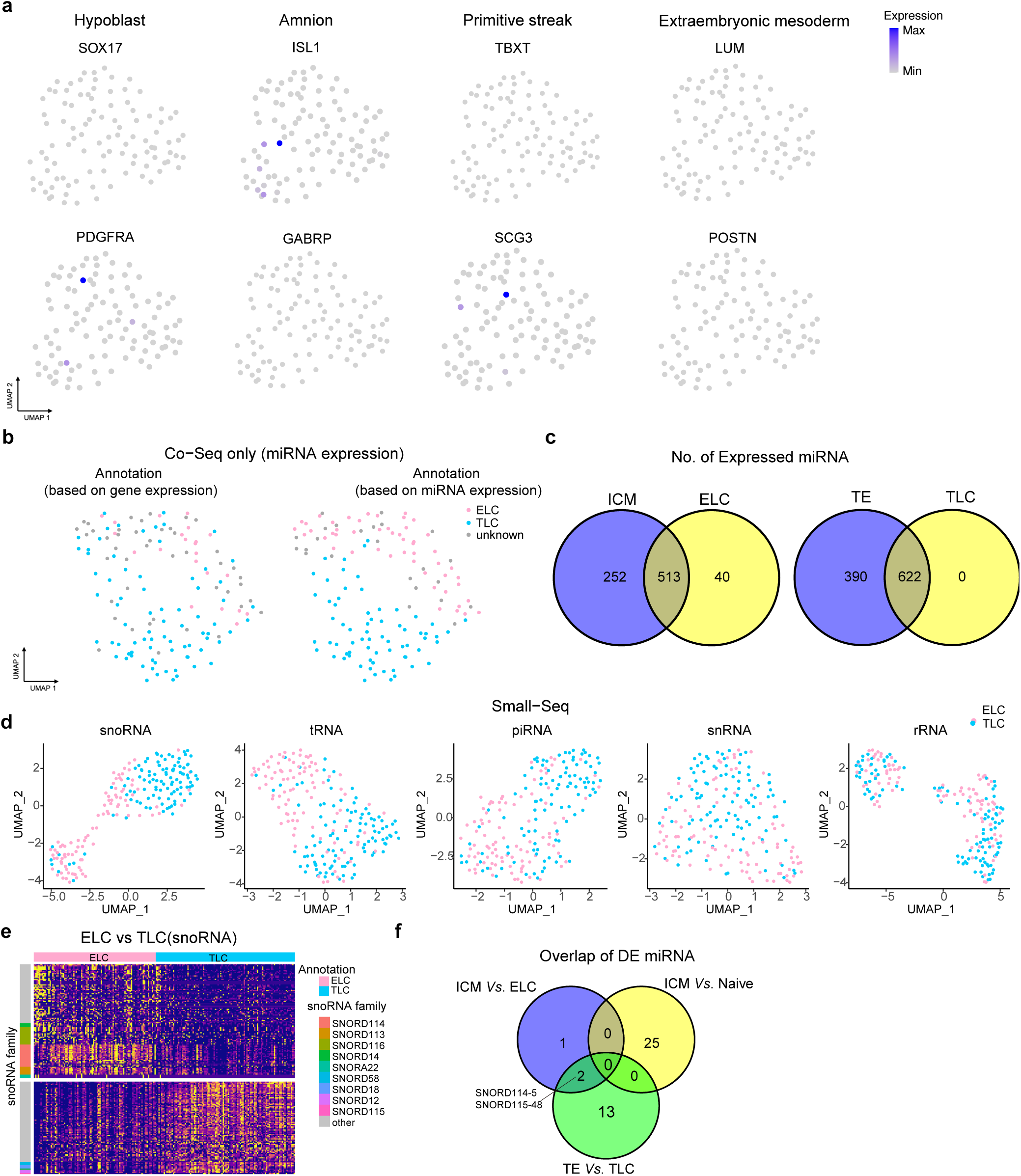
Lineage annotation and verification of human blastoid cells using mRNA and miRNA expression profiles. **a**, Expression of primitive endoderm (*SOX17*, *PDGFRA*), primitive streak (*TBXT*, *SCG3*), amnion (*ISL1*, *GABRP*), and extra-embryonic mesoderm markers (*LUM*, *POSTN*) in human blastoids Co-Seq data. **b**, UMAP by miRNA expression showing lineage annotation of human blastoids Co-Seq data (miRNA portion). **c**, Venn diagram showing the overlap of detected miRNAs between ICM and ELC, and between TE and TLC. **d**, UMAP showing dimensionality reduction results of full-cell Small-Seq blastoids data stratified by the expression of other types of sncRNAs. **e**, Heatmap displaying expression of differentially expressed (DE) snoRNA between ELC and TLC. snoRNA families with at least three members that are highly expressed in ELC or TLC are indicated by the colors on the left. **f**, Venn diagram showing the overlap of DE snoRNA among ICM vs ELC, ICM vs Naïve, and TE vs TLC comparisons.

**Extended Data Fig. 10.**
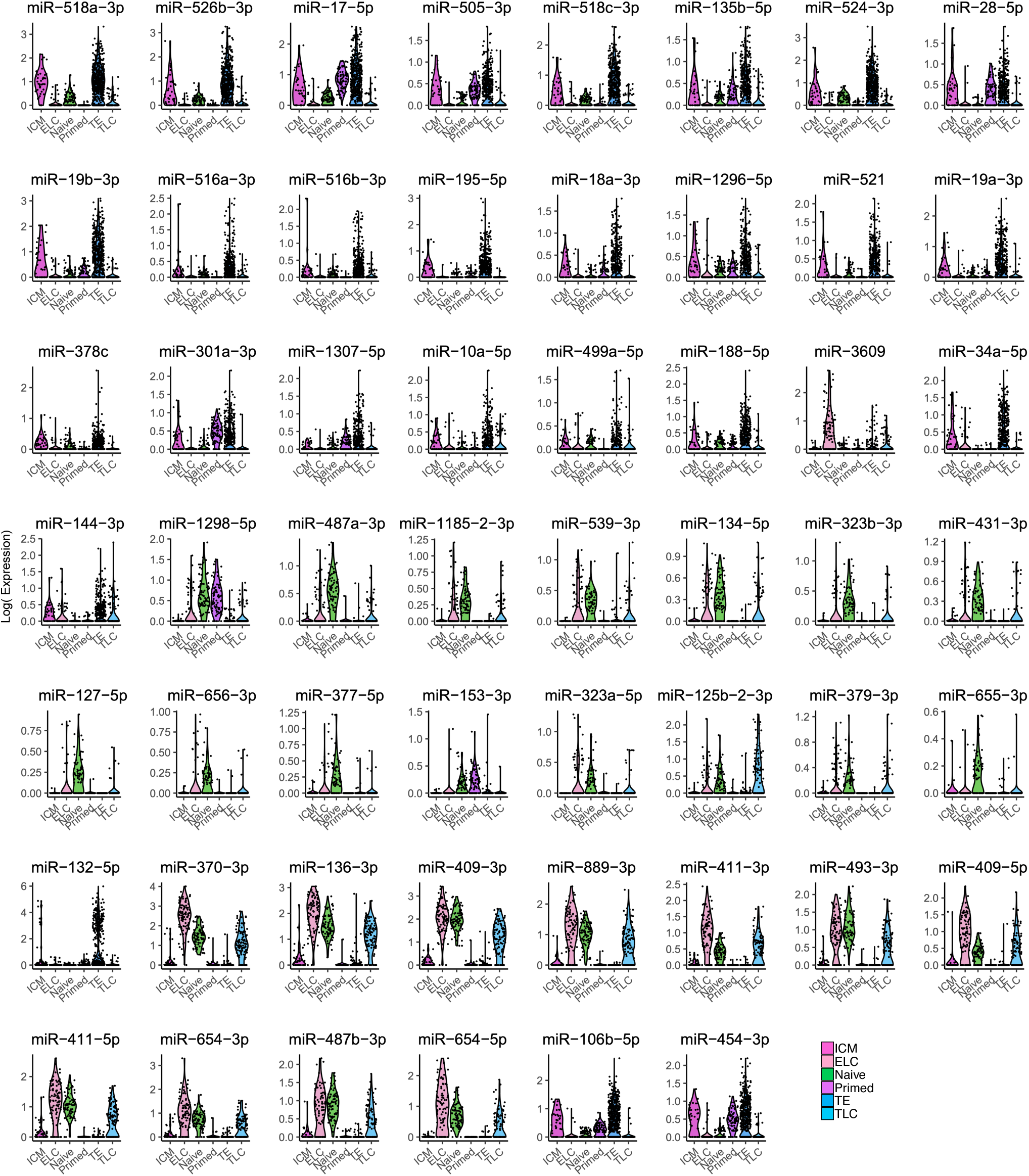
Additional miRNAs expression between human embryo, blastoid, and naive and primed human embryonic stem cells. Violin plot showing the log-normalized expression of DE miRNAs from ICM vs Naïve or ICM vs ELC or TE vs TLC across ICM, ELC, Naïve, Primed, TE, and TLC cells. ELC: Epiblast-like cells. TLC: Trophectoderm-like stem cells

